# Modeling neurodevelopmental disorder-associated *hAGO1* mutations in *C. elegans* Argonaute *ALG-1*

**DOI:** 10.1101/2023.04.06.535748

**Authors:** Ye Duan, Li Li, Ganesh Prabhakar Panzade, Amélie Piton, Anna Zinovyeva, Victor Ambros

**Author notes:** These authors contributed equally (Ye Duan, Li Li). Correspondence (A.Z.) (V.A.).

## Abstract

MicroRNAs (miRNA) are endogenous non-coding RNAs important for post-transcriptional regulation of gene expression. miRNAs associate with Argonaute proteins to bind to the 3’ UTR of target genes and confer target repression. Recently, multiple *de novo* coding variants in the human Argonaute gene *AGO1* (*hAGO1*) have been reported to cause a neurodevelopmental disorder (NDD) with intellectual disability (ID). Most of the altered amino acids are conserved between the miRNA-associated Argonautes in *H. sapiens* and *C. elegans*, suggesting the *hAGO1* mutations could disrupt evolutionarily conserved functions in the miRNA pathway. To investigate how the *hAGO1* mutations may affect miRNA biogenesis and/or functions, we genetically modeled four of the *hAGO1 de novo* variants (referred to as NDD mutations) by introducing the identical mutations to the *C. elegans hAGO1 homolog, alg-1*. This array of mutations caused distinct effects on *C. elegans* miRNA functions, miRNA populations, and downstream gene expression, indicative of profound alterations in aspects of miRNA processing and miRISC formation and/or activity. Specifically, we found that the *alg-1* NDD mutations cause allele-specific disruptions in mature miRNA profiles both in terms of overall abundances and association with mutant ALG-1. We also observed allele-specific profiles of gene expression with altered translational efficiency and/or mRNA abundance. The sets of perturbed genes include human homologs whose dysfunction is known to cause NDD. We anticipate that these cross-clade genetic studies may advance the understanding of fundamental Argonaute functions and provide insights into the conservation of miRNA-mediated post-transcriptional regulatory mechanisms.

## INTRODUCTION

The proper development, maintenance, and physiological functioning of multicellular organisms require the robust control of complex and dynamic patterns of gene expression. MicroRNAs (miRNAs) are endogenous, small, non-coding RNAs that play important roles in post-transcriptional regulation of gene expression in essentially all developmental and physiological contexts [1-3]. miRNAs are transcribed from miRNA-encoding genomic loci and undergo several processing steps to functional maturation. The initial primary miRNA (pri-miRNA) transcript is processed within the nucleus by the Microprocessor into the precursor miRNA (pre-miRNA) [2, 4]. The pre-miRNA is further processed in the cytoplasm by Dicer into the RNA duplex, which associates with a dedicated miRNA co-factor protein of the Argonaute (AGO) family. One miRNA strand from the duplex is retained by the Argonaute and becomes the functional miRNA, while the other strand is expelled from the complex and degraded. The most frequently loaded miRNA strand from the duplex is defined as the guide miRNA (miR), while the typically disposed strand is defined as the passenger (miR*). The miRNA-Argonaute complex subsequently recruits other protein factors including GW182 to form the miRNA-induced silencing complex (miRISC) [2]. In general, miRISC binds target messenger RNAs (mRNAs) at sites within in the 3’ untranslated region (3’ UTR) via partially complementary base-pairing between the miRNA and the mRNA target [2]. miRISC binding then leads to translational inhibition and/or mRNA destabilization, resulting in repression of the target gene product expression [5].

As key miRNA co-factors, the Argonaute (*AGO)* proteins are essential for miRNA-mediated post-transcriptional gene regulation [6, 7]. The *AGO* proteins participate in multiple steps of miRNA biogenesis and function, including pre-miRNA processing, miRNA duplex loading, strand selection, passenger strand disposal, target mRNA recognition, and repression of target gene expression. Accordingly, depleting *AGOs* by mutation or RNAi can result in global defects in miRNA biogenesis and target repression, and consequently lead to phenotypes characteristic of miRNA *loss-of-function* [7-11].

Interestingly, certain point mutations at conserved residues of *C. elegans* miRNA-associated Argonaute ALG-1 cause heterochronic developmental phenotypes without significant disruption of ALG-1 protein levels nor elimination of the capacity of the mutant ALG-1 protein to associate with miRNAs [12]. Strikingly, these mutants, referred to as the *alg-1* antimorphic mutants, exhibit more severe developmental defects than *alg-1 null* mutants do. Current evidence suggests that in the *alg-1 null* mutant, the loss of *alg-1* functions is largely compensated by the paralogous *alg-2* gene, whose protein product ALG-2 associates with the miRNAs that would ordinarily bind ALG-1 [7]. However, in the *alg-1* antimorphic mutants, the mutant ALG-1 antagonizes the redundancy of *alg-2* by sequestering miRNAs in defective ALG-1 miRISC, preventing those miRNAs from associating with ALG-2 [12].

*AGO* genes are implicated in multiple human diseases including male infertility, colon cancer, ovarian cancer, gastric cancer, gliomas, and neuronal developmental disorder (NDD) [13-17]. In certain cases, the disease is associated with loss of function of multiple members of the *AGO* family: Five children with psychomotor developmental delay and other non-specific neuronal-muscular disorder syndromes have been reported to be heterozygous for large *de novo* deletions in the *1p34.3* locus, which includes *hAGO1*, *hAGO3*, and *hAGO4* [18, 19]. Other cases involve *de novo* point mutations that change or delete a single amino acid of one *AGO* locus: Exome sequencing identified 18 *de novo* coding variants of *hAGO1* in children who exhibit NDD with intellectual disability (ID) and autism-spectrum disorder (ASD) [20]; similarly, 12 *de novo* coding variants of *hAGO2* were identified in children with similar spectrums of developmental delay, ID, and ASD symptoms [21]. In several cases, the same *de novo* mutations have been identified in independent families, reinforcing the conclusion that the corresponding amino acids critically contribute to AGO function. Many of the mutated amino acids are conserved between *hAGO1* and *hAGO2*, as well as between the homologous human and *C. elegans AGO* genes (Figure 1A). The conservation of these amino acids and the phenotypes associated with the corresponding mutants suggest that these amino acids are critical for evolutionarily conserved functions of AGO proteins.

**Figure 1.**
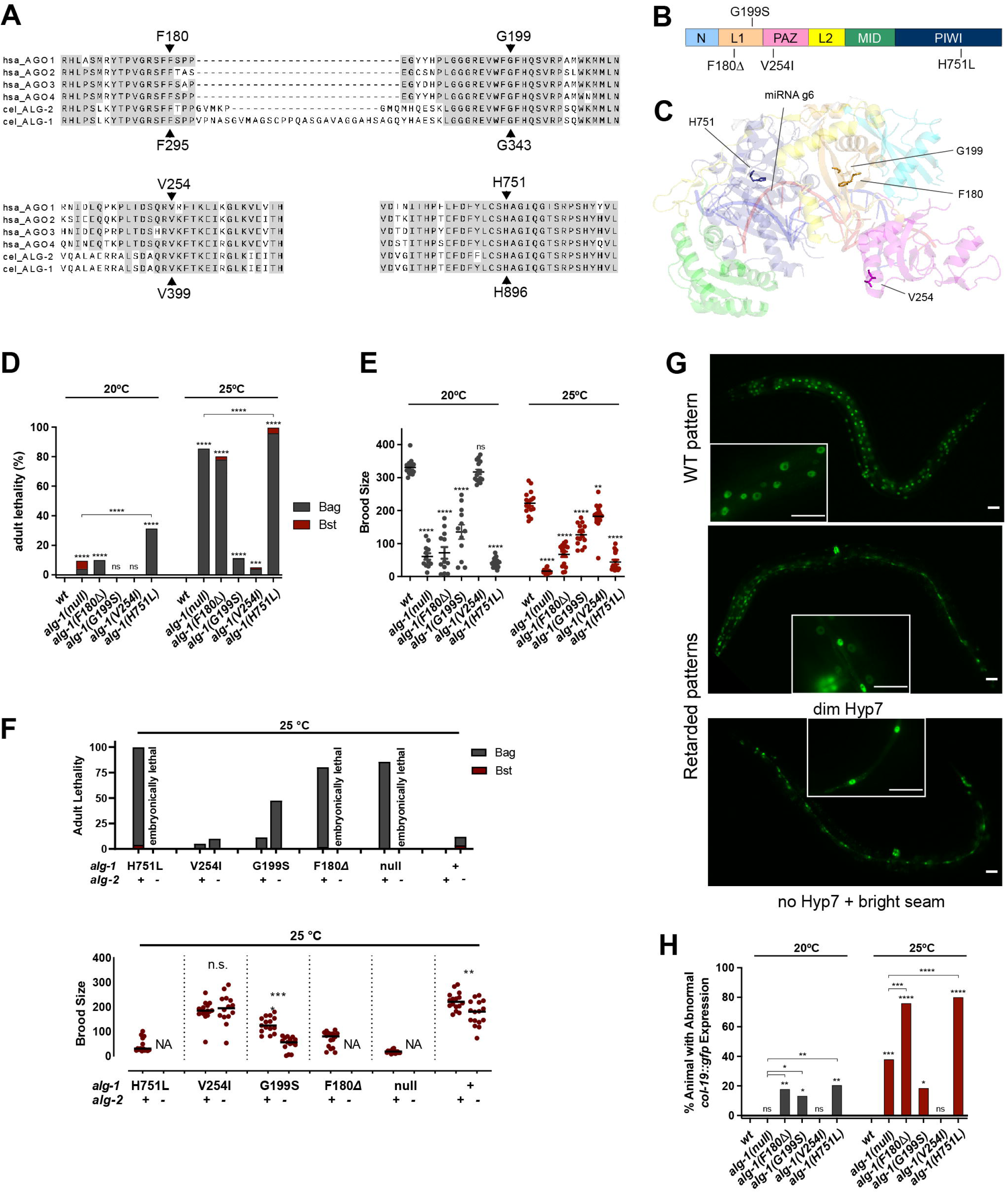
The NDD mutations cause loss-of-function and antimorphic phenotypes in the *C. elegans* Argonaute *alg-1*. **A.** Protein sequence alignment of the regions surrounding the amino acids corresponding to h*AGO1* F180, G199, V254, and H751. Alignment includes h*AGO1-4* and *C. elegans* ALG-1 and ALG-2. The ALG-1 amino acid numbers (indicated at the bottom) correspond to *C. elegans* ALG-1 isoform a (ALG-1a). Alignment is analyzed by CLUSTALW [76]. **B.** Domain organization of *C. elegans* ALG-1. The unstructured and non-conserved sequence at the N-terminus (aa 1-187) of cel-ALG-1a is not shown. **C.** hAGO2::miRNA::target complex structure (PDB:: 6MFR) with the localization of hAGO1 F180, G199, V254, H751 residues [31]. Side chains of the above amino acids are presented as sticks. **D-E**. Quantification of vulval defect phenotypes, represented by the lethality of young adult hermaphrodites (**D**) and reduction in the number of progeny per animal (**E**). The lethality is categorized as due to vulval integrity defect (lethality by bursting, Bst) or egg laying defect (lethality by matricide, wherein embryos hatch within and consume the mother, Bag). The vulval integrity defect (Bst) is considered the more severe phenotype. **F.** Quantification of vulva integrity defect (left) and abnormal *col-19::gfp* expression defect (right) of the *alg-1* NDD mutations with *alg-2(+)* or *alg-2(null)* genetic backgrounds. **G.** Representative fluorescent images of *col-19::gfp* expression patterns for the phenotypic scoring. Images showing WT *col-19::gfp* expression pattern are taken from *maIs105.* Images showing dim Hyp7 expression patterns are taken from *maIs105; alg-1(G199S).* Images representing no Hyp7 expression pattern is taken from *maIs105; alg-1(H751L).* Scale bar is 25 µm. **H.** Quantification of the *col-19::gfp* expression defect phenotypes. The statistical significance of lethality and abnormal *col-19::gfp* expression are analyzed by Fisher’s test. The statistical significance of brood size is analyzed by Student t-test (see Method). ****p≤0.0001, ***p≤0.001, **p≤0.01, *p≤0.05.

It is noteworthy that among the described NDD cases, frameshift mutations or large deletions that could result in unambiguous single-gene *null* mutations of *hAGO1* or *hAGO2* were rarely documented [22]. This suggests that the NDD-associated single amino acid mutations of *hAGO1* or *hAGO2* may be more malicious than either *null* allele, perhaps by antagonizing otherwise redundant paralogous *AGO* genes. Interestingly, two of the NDD-related *de novo* mutations (corresponding to H751L and C749Y in *hAGO1*) are adjacent to the previously described antimorphic allele (corresponding to S750F in *hAGO1*) in *C. elegans alg-1* (Figure 1A) [12]. Thus, it is likely that some of the NDD-related *AGO* mutations may have antimorphic impact on *AGO* function. We thus reasoned that modeling the NDD-related human *AGO* mutations in conserved *C. elegans AGO* mutations could provide a rapid way to assess the effects of the mutation on miRNA biogenesis and miRNA/AGO functionality.

Here, we reproduced four *hAGO1* mutations (F180Δ, G199S, V254I, H751L) in the homologous *C. elegans alg-1* gene using CRISPR/Cas9-mediated genome editing. We refer to the corresponding *C. elegans* mutations as *alg-1* NDD mutations. We show that the *alg-1* NDD mutations resulted in developmental phenotypes ranging from loss-of-function to antimorphic, with two *alg-1* NDD mutations resulting in stronger heterochronic phenotypes than the homozygous *alg-1 null* mutants. The antimorphic character of *alg-1* NDD mutations suggests that the mutant ALG-1 protein interferes or competes with the functions of paralogous Argonaute proteins (nominally ALG-2 in *C. elegans*). We found that *alg-1* NDD mutations affected the overall profile of mature miRNAs and the profile of miRNAs associated with ALG-1 protein, including the proper selection of mature miRNA guide/passenger strands. We also observed that the mutations caused global gene expression perturbations in terms of both mRNA levels and mRNA translational status, including substantial differences in the de-repression modes of miRNA targets for certain mutations. Interestingly, the set of *alg-1* NDD mutations examined here exhibit distinguishable allele-specific perturbations in *C. elegans* miRNA function, miRNA profiles, and gene expression, suggesting that the NDD mutations each impair ALG-1 functionality with allele-specificity. Lastly, we show that a large proportion of the genes whose expression is perturbed by the *alg-1* NDD mutations are known to have human homologs whose dysfunction is known to cause NDD. Our results demonstrate that modeling *hAGO1* mutations in a *C. elegans* Argonaute can advance the understanding of fundamental Argonaute functions and provide insights into the conservation of miRNA-mediated regulatory mechanisms.

## RESULTS

### The NDD mutations disrupt ALG-1 Argonaute function

We selected four *hAGO1* NDD-related mutations (F180Δ, G199S, V254I, H751L) to model in *C. elegans* ALG-1. All four mutations had been identified in multiple patients, with the F180Δ, G199S, and V254I mutations identified in independent families [20]. Further, the F180Δ and G199S mutations were also identified at homologous positions of *hAGO2,* causing NDD with ID and ASD symptoms [21]. The amino acids mutated in *hAGO1* (F180, G199, V254, H751) are conserved among the four miRNA-associated *AGO* proteins in humans (*hAGO1*-*hAGO4*), as well as their two *C. elegans* orthologs, ALG-1 and ALG-2 (Figure 1A). The mutated F180 and G199 amino acids reside in the L1 hinge domain, which lies between the MID-PIWI lobe and the PAZ-N lobe (Figure 1B-C). The V254 amino acid resides in the PAZ domain with the side-chain exposed to the surface of AGO1 protein and is distant from the PAZ-N channel where the 3’ non-seed duplex is located (Figure 1B-C). The H751 amino acid resides in the PIWI domain and is near the MID-PIWI channel where the miRNA seed duplexes with the target (Figure 1B, 2A).

**Figure 2.**
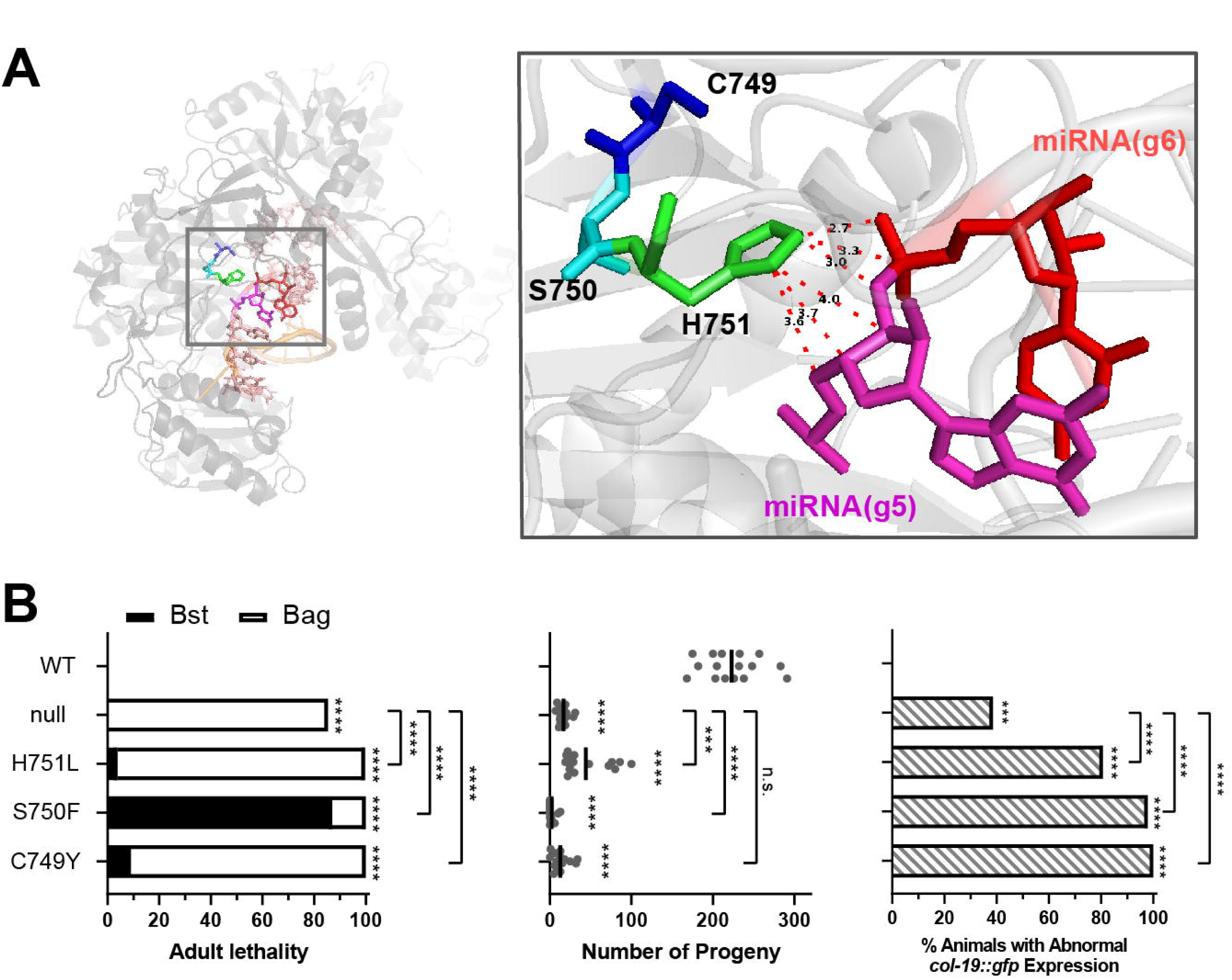
The C749-S750-H751 subregion is functionally critical to ALG-1. **A.** Visualization of the hAGO2 side-chains equivalent to hAGO1 C749, S750, and H751 and g5-g6 nucleotides of miRNA in the AGO2::miRNA::target complex (PDB:: 6MDZ) [31]. Dashed lines and numbers indicate distances between adjacent atoms (Å). **B.** Lethality, brood size, and abnormal *col-19::gfp* expression phenotypes of the *alg-1(C749Y)*, *alg-1(S750F)*, and *alg-1(H751L)* mutants. Phenotypes are scored at 25 °C. ****p ≤ 0.0001. ***p ≤ 0. 001.

To explore how the NDD mutations may affect the functionality of miRNA regulation, we used CRISPR/Cas9 genome editing to generate four *C. elegans* mutant strains, each containing a mutation identical to F180Δ, G199S, V254I, or H751L at the corresponding amino acids of *C. elegans* ALG-1. Note that in this paper, we refer to each *C. elegans* ALG-1 mutation using the human *AGO1* addresses of the corresponding amino acid (Figure 1A). The respective *C. elegans* mutants are *alg-1(ma447, F180Δ*), *alg-1(ma443, G199S), alg-1(zen25, V254I)* and *alg-1(zen18, H751L)*.

*C. elegans* animals homozygous for each of the four *alg-1* NDD mutations exhibit varying degrees of developmental defects (Figure 1D-H). *alg-1(*F180Δ) and *alg-1(*H751L) hermaphrodites exhibit strong adult lethality (Figure 1D), caused by impaired egg laying (retention of embryos and eventual matricide by the progeny that hatch *in utero*) and/or rupturing of the cuticle at the vulva, which kills the adult outright before reproduction. Both these phenotypes are presumed to reflect defective vulval development owing to decreased activity of certain miRNAs known to be critical for normal vulva development [7]. In accordance with their underlying egg retention and/or vulva bursting defects, *alg-1* NDD mutant hermaphrodites produced a dramatically reduced number of progeny (Figure 1E). The penetrance of the adult lethality and reduced progeny phenotypes for the H751L and F180Δ mutants were at least as strong as for *alg-1(tm492, null)* mutants (Figure 1D-E). *alg-1(G199S)* animals exhibited moderate penetrance of vulval defects and reduced number of progeny, and *alg-1(V254I)* mutants exhibited relatively mild and temperature-dependent expression of these phenotypes (Figure 1D-E). Thus, based on phenotypic comparison with *alg-1 null,* we conclude that the F180Δ, G199S, V254I, and H751L mutations cause varying degrees of ALG-1 loss-of-function [7, 12].

### The *alg-1* NDD mutations synergize with the *alg-2 null*

Functional redundancy has been reported among miRNA-related *AGO* genes, both for the human Ago family (*hAGO1*-h*AGO4*) and for the *C. elegans alg-1* and *alg-*2 [7, 12, 23]. To test for redundancy associated with the functions disrupted by the *alg-1* NDD mutations, we crossed each of the four *alg-1* mutations into the *alg-2(ok304, null)* genetic background. We found that the *alg-1(F180Δ*); *alg-2(null)* and *alg-1(H751); alg-2(null)* mutants exhibited embryonic lethality, consistent with severe reduction of both *alg-1* and *alg-2* functions [7] (Figure 1F). Meanwhile, the weaker *alg-1* NDD mutations also exhibited genetic interactions with *alg-2 null* mutation: *alg-1(G199S); alg-2(null)* mutants showed increased adult lethality and reduced number of progeny compared to *alg-1(G199S)* and *alg-1(V254I);alg-2(null)* animals exhibited increased adult lethality compared to *alg-1(V254I)* (Figure 1F). Since both the vulva developmental defect and the reduction in the progeny of all four *alg-1* NDD mutations were exacerbated in the *alg-2 null* background, we conclude that the *alg-1* NDD mutations disrupt, to varying degrees, *alg-1* functions that are redundant with *alg-2*.

### *alg-1(F180Δ)* and *alg-1(H751L)* mutations are antimorphic in regulating seam cell differentiation

In *C. elegans,* lateral hypodermal development involves a stem cell lineage wherein the stem cells (seam cells) execute asymmetric divisions at each larval stage, producing one daughter cell that differentiates and joins the hypodermal syncytium (Hyp7) and another daughter cell that remains a stem cell [24]. At the final larval molt, seam cells cease division, and all hypodermal cells (seam and Hyp7) express the adult-specific hypodermal gene *col-19* [25, 26]. miRNAs – particularly the *lin-4* family and *let-7* family miRNAs --are critical for controlling the timing of this larval-to-adult hypodermal cell fate transition [3, 27, 28]. Accordingly, mutations that disrupt these miRNAs or the miRNA machinery can cause the failure of hypodermal cells to properly express adult fates, as reported by the expression of the *col-19::gfp* transgene, including reduced or absent *col-19::gfp* expression in Hyp7 [12, 29] (Figure 1G).

We found that *alg-1(F180Δ)*, *alg-1(H751L),* and *alg-1(G199S)* adults exhibited reduced or absent *col-19::gfp* expression in Hyp7 cells, indicating that the NDD mutations impair heterochronic pathway miRNA activity (Figure 1H). Strikingly, the heterochronic phenotypes of the *alg-1(H751)* (80.7%) and *alg-1(F180Δ)* (76.6%) were significantly stronger than that of the *alg-1(null)* mutant (38.6%) (Figure 1H). Since *alg-1(H751L)* and *alg-1(F180Δ)* homozygotes exhibit phenotypes stronger than *alg-1(null)* homozygotes, we conclude that the *alg-1(F180Δ)* and *alg-1(H751L)* mutations are antimorphic and that these mutations do not simply inactivate the ALG-1 protein, but alter ALG-1 function such that the mutant protein inappropriately interferes with the function of other gene products – in this case presumably ALG-2. The antimorphic behavior of the *alg-1(F180Δ)* and *alg-1(H751L)* mutations is reminiscent of other *C. elegans alg-1* alleles that were identified in forward screens for heterochronic mutants and are likewise point mutations at evolutionarily conserved amino acids [12].

Interestingly, in animals carrying an *alg-1* NDD mutation heterozygous to a wild-type *alg-1* allele, no lethality or abnormal *col-19::gfp* expression was observed (Figure S1A-B). Meanwhile, the reduction of the number of progeny of the heterozygous mutants, if any, was not as remarkable as for the corresponding homozygotes (Figure S1C). This indicates that the *alg-1* NDD mutations, similar to the *alg-1* antimorphic mutations described previously, appear to be fully recessive or only weakly semi-dominant [12]. Thus, the *alg-1* NDD mutations are not strictly dominant negative in the classical sense, and their negative activities may be dosage dependent or complemented by a WT allele.

### The C749-S750-H751 sub-region is critical for ALG-1 function

One of the previously-described *C. elegans alg-1* antimorphic mutations, *alg-1(ma192),* is a serine-to-phenylalanine point mutation at the amino acid homologous to S750 of *AGO1*, which is adjacent to the H751L NDD mutation. In addition, another human genetic study of NDD patients reported three independent cases of a cysteine-to-tyrosine change in *hAGO2* at the amino acid homologous to C749 of *hAGO1* [21]. It is striking that three independent genetic screens in human and *C. elegans* have recovered mutations at three adjacent Argonaute amino acids (C749Y, S750F, and H751L) that seem to cause a particularly potent class of defects. The adjacency of the three mutations suggests that these amino acids lie in a region of Argonaute protein critical for function and may affect the activity of the protein similarly.

The current structure of co-crystalized hAGO2::miRNA::target ternary complex supports the hypothesis that the C749-S750-H751 region could be particularly critical for Argonaute function (Figure 2A) [30, 31]. The PIWI domain amino acids affected by the identified mutations are positioned close to the backbone of the miRNA at g5-g6 seed nucleotides (Figure 2A). Particularly, most atoms in the imidazole group of the H751 side chain are spatially close to the atoms of the backbones of miRNA g5 and g6 nucleotides, with an average distance of 3.6 Å, suggesting that the side-chain of H751 directly contacts the miRNA seed region via hydrogen bonds and electrostatic force [32] (Figure 2A).

To further investigate and compare the functions affected by the C749, S750, and H751 mutations, we introduced the C749Y mutation into the *C. elegans alg-1* by CRISPR/Cas9, and compared its phenotypes to those of *alg-1(S750F)* and *alg-1(H751L)* mutants. Like S750F and H751L, *alg-1(ma545, C749Y)* mutant animals exhibited adult lethality, reduced number of progeny, and abnormal *col-19::gfp* expression (Figure 2B). Moreover, all three mutants exhibited a penetrance of the *col-19::gfp* expression defect that is greater than the *alg-1(null)* mutant, suggesting that all these three mutations confer antimorphic activity to ALG-1 (Figure 2B). These findings support the conclusion that the C749-S750-H751 sub-region is particularly critical for certain Argonaute functions, such that mutations in this region can cause the mutant ALG-1 protein to antagonize the functions of ALG-2.

### The *alg-1* NDD mutations can disrupt total miRNA profiles and the profiles of miRNAs associated with ALG-1

Previous studies of *C. elegans alg-1* antimorphic mutations reported global abnormalities in miRNA biogenesis without affecting Argonaute protein levels [12, 29]. The expression levels of the mutated ALG-1 NDD proteins similarly did not differ significantly from WT ALG-1, suggesting that overall protein stability was not affected by these mutations (Figure 3). We then sought to test whether the *alg-1* NDD mutations can cause similar perturbations in the expression of *C. elegans* miRNAs. We performed small RNA sequencing (sRNA-seq) of total RNA from L4 larval extracts and analyzed the expression levels of guide strands for the 259 relatively abundant (minimal RPM > 5) miRNAs (Figure 3A-B, S2, S3, Table S1). We found that the F180Δ and G199S mutations caused a remarkable disturbance of total miRNA profiles, with 87 and 75 miRNAs, respectively, perturbed more than 2-fold (FDR < 0.05) (Figure 3C-D). Only one miRNA was changed in level with statistical significance in the V254I mutant (Figure S3), which is consistent with the weak phenotypes of *alg-1(V254I)* (Figure 1D-H). Surprisingly, only 21 miRNAs were significantly perturbed in the H751L mutants (Figure 3C-D), which is in sharp contrast with the strong phenotypes of H751L mutant animals.

**Figure 3.**
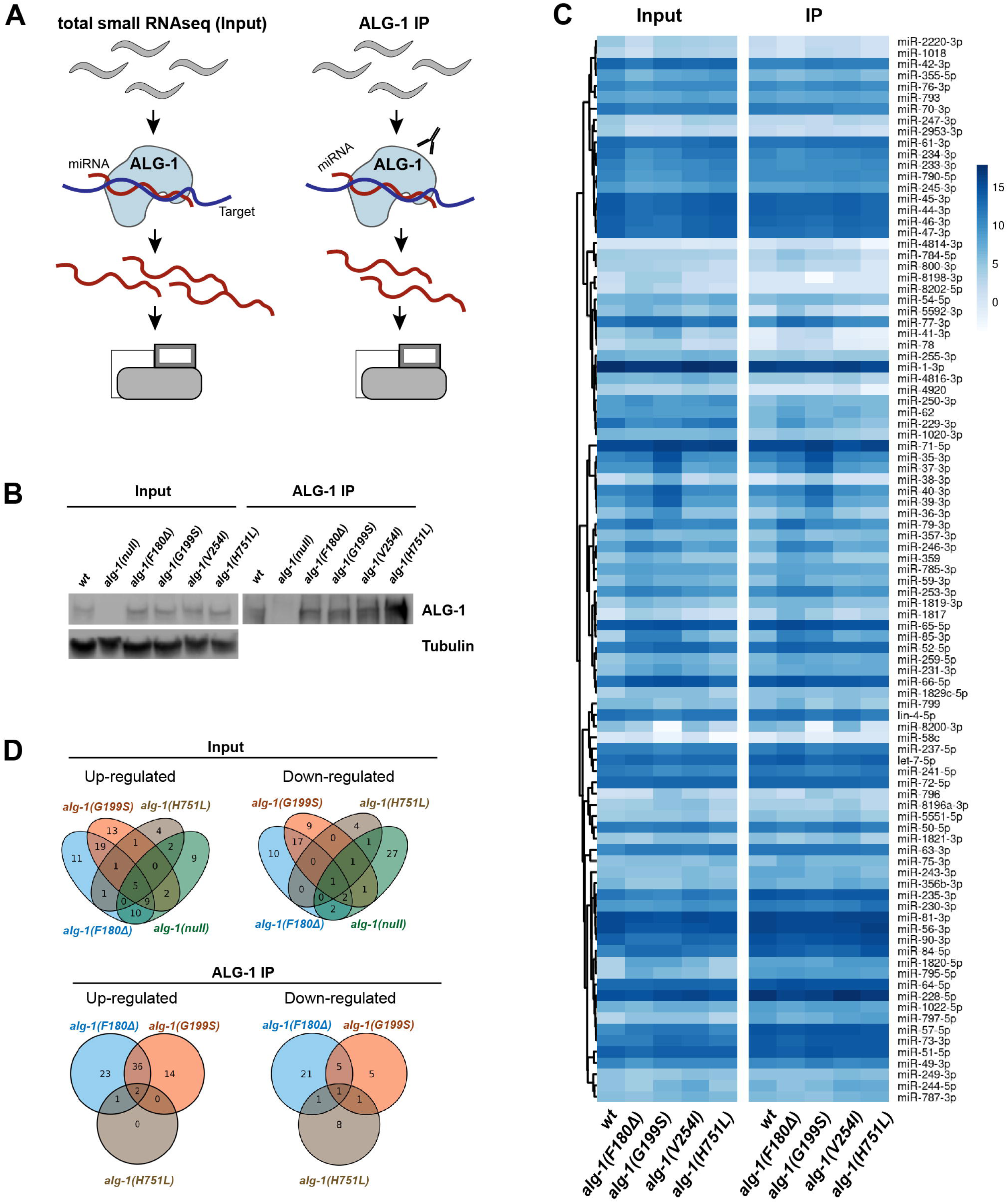
The *alg-1(NDD)* mutations cause allele-specific disruptions of miRNA expression and miRNA associated with ALG-1. **A.** Schematic diagram of ALG-1 IP and small RNA sequencing (input). **B.** Western-blotting for ALG-1 protein in input and ALG-1 immunoprecipitated samples of wild type, *alg-1(null),* and *alg-1(NDD)* mutants. Tubulin is detected as the loading control for input samples. **C.** Heatmap showing the levels of abundant miRNAs (≥10rpm) in the input and ALG-1 IP samples of wild type and *alg-1(NDD*) mutants. Data are shown as log2(RPM). **D.** Venn diagrams showing numbers of miRNAs with statistically significant up/down-regulated levels (Fold change > 2 and FDR < 0.05) in the input (top) and ALG-1 IP (bottom). Results for *alg-1(V254I)* mutant are not shown because no significant perturbation was observed in ALG-1(IP), while only a single miRNA was up-regulated in *alg-1(V254I)* input (Figure S3 and Table S1).

To determine whether the *alg-1* NDD mutations affect the profiles of miRNAs that associate with the mutant ALG-1 protein, we performed ALG-1 immuno-precipitation (IP) using anti-ALG-1 polyclonal antibody and sequenced the miRNAs co-immuno-precipitated with ALG-1 (Figure 3A-B, S2A). Consistent with the observed changes in total miRNAs (input), many miRNAs showed altered association with mutant ALG-1 compared to the wild-type (Figure 3C-D, S2C-D, S3, Table S1). Specifically, among the miRNAs that co-immuno-precipitated with ALG-1, 90 miRNAs for F180Δ, 64 miRNAs for G199S, 14 miRNAs for H751L, and 0 miRNAs for V254I were significantly changed compared to those co-immunoprecipitated with wild-type ALG-1 (Figure 3C-D). The majority of the perturbed miRNAs that exhibited altered expression in the IP also had a corresponding change in the input with statistically significant enrichment (Figure S2A).

It is noteworthy that while some miRNAs were perturbed in multiple NDD mutants, some miRNAs were uniquely perturbed in individual mutants (Figure 3D). This observation underscores the suggestion that different NDD mutations affect ALG-1 function differently, possibly reflecting the unique functions of the affected protein domains (Figure 3D). In addition, large proportions of the miRNAs disrupted in the NDD mutants were distinct from the miRNAs affected in the *alg-1 null* (Figure S2B), suggesting that the miRNA perturbations in the NDD mutants are not simply caused by a reduction of *alg-1’s* normal contribution to miRNA biogenesis.

### The NDD mutations can alter guide/passenger strand ratios

In the previous report, *C. elegans alg-1* antimorphic mutations, including the *alg-1(S750F)* mutation which is adjacent to H751L, disturbed miRNA biogenesis not only by changing miRNA expression levels, but also by altering relative abundances of guide (miR) and passenger strands (miR*) for particular miRNAs [12, 29] (Figure 4A). We therefore asked whether the *alg-1* NDD mutations can similarly cause alterations in relative miR/miR* strand abundance. We found that, for multiple miRNAs, the miR*/miR ratios were altered in the F180Δ, G199S and H751L mutants (Figure 4B-D). In principle, an altered miR/miR* ratio can reflect altered guide-passenger selection as discussed above, or defects that do not alter strand choice *per se*, but affect the relative stability of the two strands, for example by a failure to dispose the passenger strand. To distinguish these two scenarios, we examined the expression levels of guide and passenger strands of individual miRNAs in the NDD mutants (Figure 4E). We observed that some of individual miRNAs exhibited increased expression of miR* accompanied by decreased expression of the guide strands in the NDD mutants, suggesting that the changed miR/miR* ratio could largely be attributed to a strand selection defect (Figure 4E).

**Figure 4.**
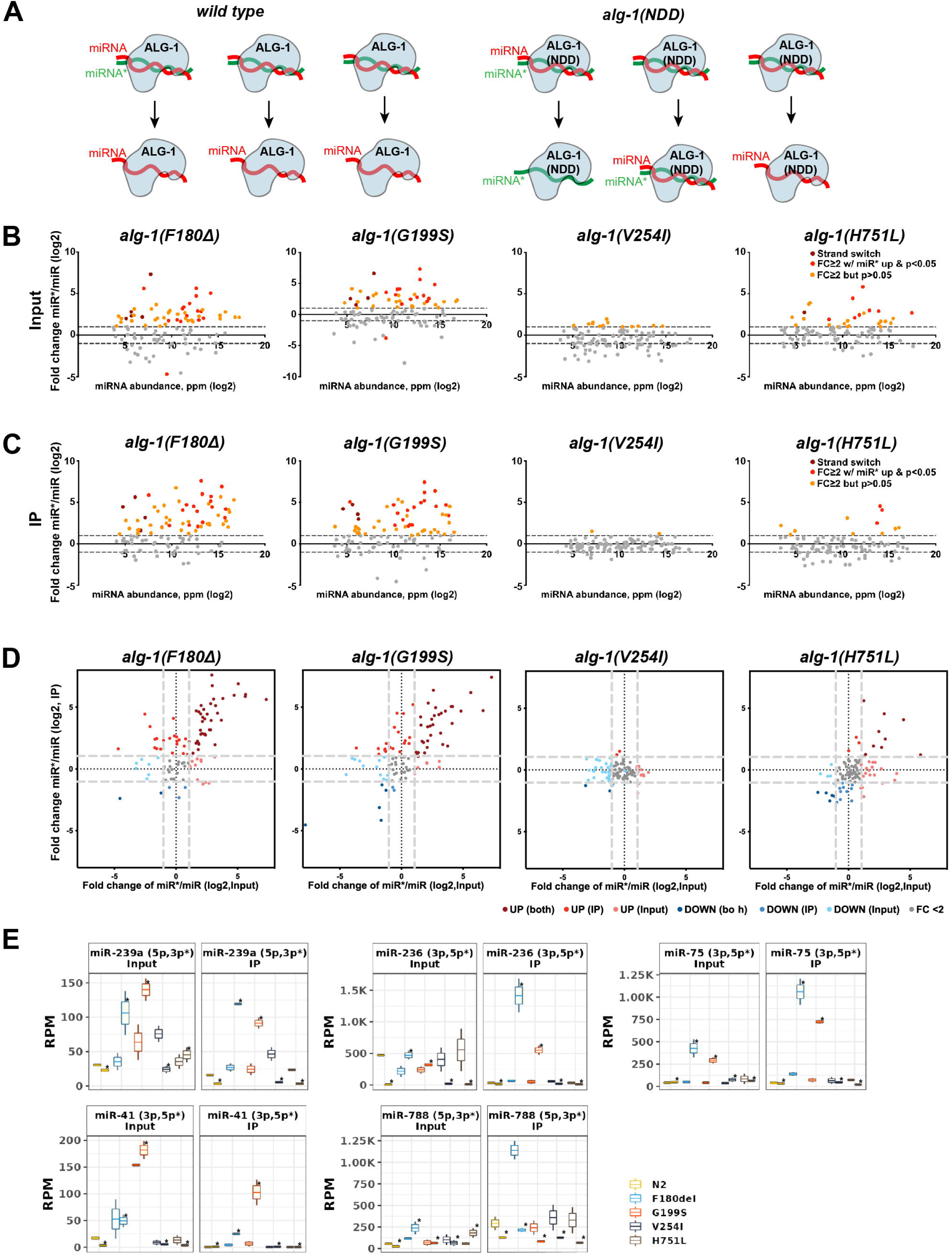
*alg-1* NDD mutations lead to altered guide/passenger (miR/miR*) ratios. **A.** A schematic model of ALG-1 miRNA strand loading in wild type and NDD mutant ALG-1. **B-C.** Changes of miR/miR* ratio in the input (B) and ALG-1 IP (C). log2FC miR*/miR ratio in wild type vs. mutant animals (Y-axis) is plotted against miRNA abundance in wild type (X-axis). Burgundy dots represent miRNAs with switched miRNA strand abundance (miR*>miR), red dots represent miRNAs whose miR* strands were upregulated ≥2 fold with p≤0.05, and orange dots represent miRNAs whose miR* strands were upregulated ≥2 fold but did not reach statistical significance. Dashed lines, |log2FC| = 1. **D.** miRNA fold change comparison between input and ALG-1 IP. miRNAs with |FC|>2 and p < 0.05 are color-coded to indicate miRNA up-or down-regulation in both input and ALG-1 IP, and input or ALG-1 IP only. **E.** miRNAs that exhibited reversed miRNA strand abundance in input and/or ALG-1 IP. miRNA* strands are marked with an asterisk(*).

### The *alg-1* NDD mutations cause allele-specific translatome-wide perturbations in gene expression

The AGO protein is a core miRISC component and therefore is critical for post-transcriptional gene regulation. Changes in ALG-1 functions could be expected to disrupt gene expression profiles due to the de-repression of protein production from mRNAs directly targeted by miRNAs, combined with indirect perturbation of genes downstream of disrupted miRNA targets. To assess how the NDD mutations affect genome-wide gene expression as changes in translation from each mRNA, we used ribosome profiling (Ribo-seq) to profile ribosome occupancy of mRNAs in extracts of late L4 animals for the WT, *null*, and NDD mutants [33, 34]. We observed that all the mutations can perturb the translatome of the mutant animals compared to the WT (Figure 5A, Table S2). The number of genes with statistically significant perturbations in ribosome protected fragment (RPF) counts (|FC|>2 and p.adj < 0.1) ranged from 66 genes (V254I) to 1731 genes (F180Δ) (Figure 5A). The gene expression changes were observed for both abundantly expressed genes and genes with low expression levels (Figure S4F). PCA analysis suggests that each NDD mutant exhibits a distinctively perturbed translatome (Figure 5B), and the sets of genes perturbed in the weaker mutants (i.e., V254I) were not simply a subset of the sets perturbed in the stronger mutants (Figure 5C), suggesting that each mutation may impair ALG-1 function in a qualitatively distinct fashion.

**Figure 5.**
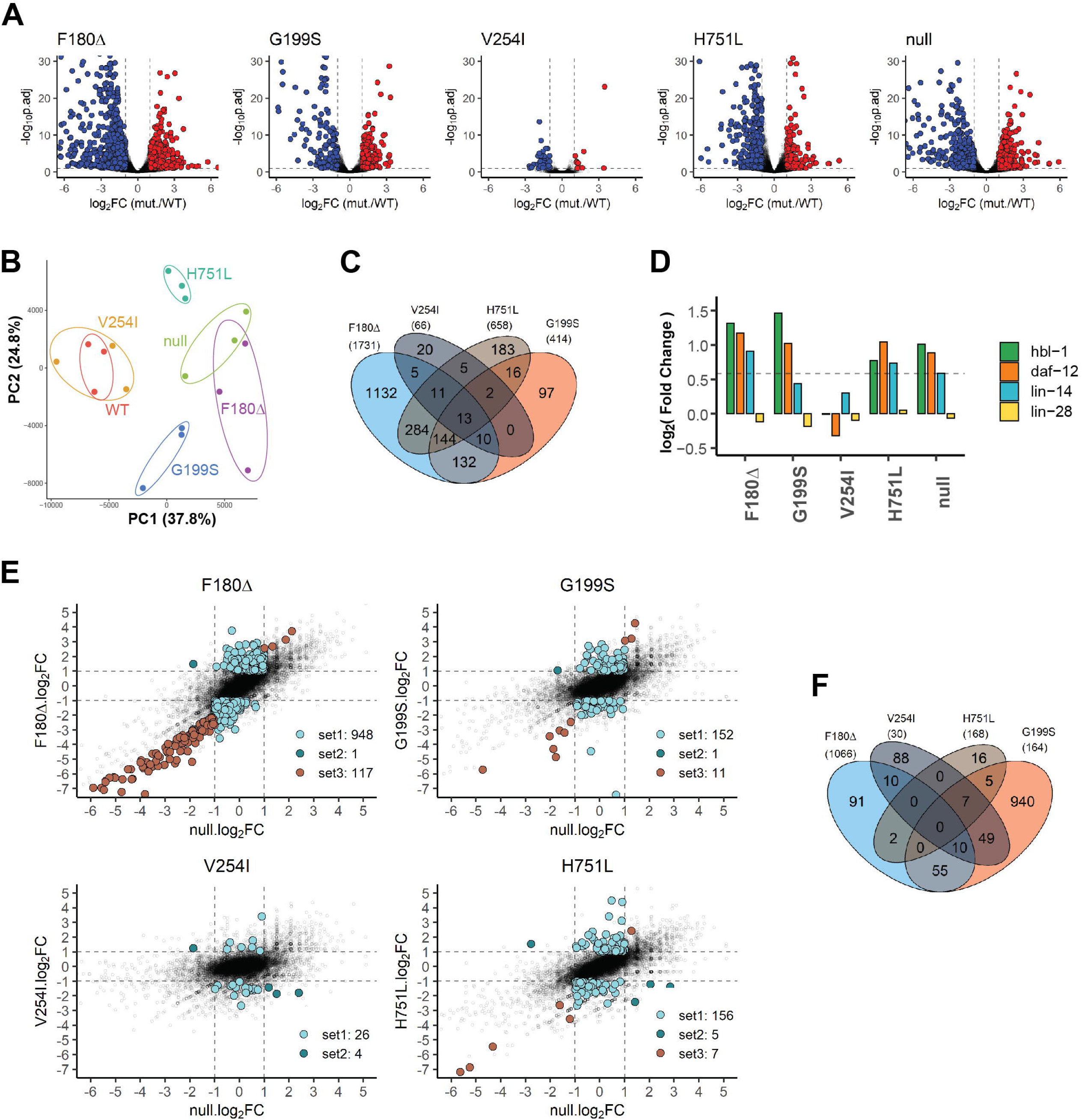
*alg-1* NDD mutations lead to strong translatome perturbations in *C. elegans*. **A.** Volcano plots of the ribosome protected fragments (RPF) detected in ribosome profiling of NDD mutant late L4 larvae. Colored dots represent perturbed genes with statistical significance (|log2FC| > 1.5, p.adj < 0.1). Also see Table S2. **B.** Principal component analysis plot of translatomes of the NDD mutants and *alg-1 null*. Points with identical colors indicate biological replication. **C.** Venn diagram for the total perturbed genes in the NDD mutants. **D.** RPFs of heterochronic genes whose gain-of-function mutations were reported to cause heterochronic phenotypes and have been genetically confirmed to be miRNA targets in *C. elegans*. **E.** Visualization of set1-set3 <u>a</u>nti<u>m</u>orphic <u>p</u>erturbed (*amp*) genes. For each gene, the log_2_FC of the *null*/WT is plotted on the x-axis and the log_2_FC of NDD mutant/WT is plotted on the y-axis. Solid dots indicate perturbed genes with statistical significance ( |FC| >1.5, p.adj < 0.1). See also Table S3. **F.** Venn diagrams of set1-set3 *amp* genes in the NDD mutants.

### The major heterochronic genes were translationally perturbed in NDD mutants

The *alg-1* NDD mutants exhibit adult lethality and *col-19::gfp* expression defects in the hypodermis, consistent with disruptions in miRNA-regulated heterochronic pathway function. We found that genes that were translationally up-regulated in *alg-1* NDD mutants are enriched for genes expressed in hypodermal seam cells (Figure S4E). The enriched seam cell genes included major heterochronic genes *daf-12*, *hbl-1*, and *lin-14* (Figure 5D). Gain-of-function mutations in these genes have been reported to cause heterochronic phenotypes, and these genes have been genetically confirmed to be miRNA targets [3, 26, 35-38]. We found that the translation of *daf-12* and *hbl-1* was up-regulated in the F180Δ, G199S, and H751L mutants, and *lin-14* was up-regulated in the F180Δ and H751L mutants (Figure 5D). This observation suggests that the abnormal function of ALG-1 NDD miRISC causes over-expression of these heterochronic genes and consequently leads to the developmental phenotypes in the *alg-1* NDD mutants. Meanwhile, none of the major heterochronic genes were significantly perturbed in the V254I mutant, consistent with the mild phenotypes of V254I animals.

### The NDD mutations cause antimorphic translatome perturbations

Consistent with *alg-1* NDD mutations causing *alg-1* loss-of-function, the translationally perturbed genes in *alg-1* NDD mutants partially overlap with the genes perturbed in the *alg-1 null* mutants (Figure S4A). However, the translatome perturbations of the NDD mutants are also strikingly distinct from *alg-1 null* mutant because they include gene changes not observed in the *alg-1(null)* (Figure 5E, set1, Figure S4B). In addition, some genes were found up-regulated in *alg-1* NDD mutants but were down-regulated in the *alg-1 null* mutant (Figure 5E, set2, S4C). Other genes were perturbed in both the *alg-1* NDD mutants and *alg-1 null* mutants but with greater perturbation (|ΔFC|> 2) in the *alg-1* NDD mutants than in the *null* mutant (Figure 5E, set3, S4D). Together, these genes form a gene subset distinctly affected in *alg-1* NDD animals versus *alg-1 null* animals. We refer to these genes (set1-3) as <u>a</u>nti<u>m</u>orphic <u>p</u>erturbed (*amp*) genes (Table S3). For the F180Δ, G199S, and H751L mutations, the occurrence of the *amp* gene perturbations in excess of those in *alg-1 null* is consistent with the observation that these mutations cause developmental phenotypes stronger than *alg-1 null* (Fig. 1D-H). Interestingly, even the weakest mutation V254I, which displayed negligible visible phenotypes, nevertheless exhibited *amp* genes of the set1 and set2 classes (Figure 5E). We propose that the perturbation of the *amp* genes may contribute to the antimorphic phenotypes of *alg-1* NDD mutants and that all the NDD mutations that we modeled in *C. elegans* confer an antimorphic impact on the ALG-1 protein in the context of translatome regulation.

### The translatome disruptions in *alg-1* NDD mutants correspond to distinct profiles of miRNA perturbation

The relatively mild perturbation of miRNA levels in the total miRNA and ALG-1 IP profiles caused by the H751L mutation stands in striking contrast with the severity of the developmental phenotypes, as well as the substantial perturbations in the translatome exhibited by H751L mutants. Specifically, the H751L mutation causes stronger developmental phenotypes and gene perturbation than G199S (Figure 1D-F, Figure 5A-B). However, the number of miRNAs whose total abundance or ALG-1 association is reduced in the H751L mutant are far less than the number of miRNAs reduced in the G199S mutant (Figure 3C-D). This contrast suggests that the stronger phenotypes of H751L mutant animals reflect a substantial loss of miRNA function that is not reflected by miRNA levels. We therefore hypothesized that the H751L mutant ALG-1 protein, although relatively normal for miRNA biogenesis/association, is defective in one or more subsequent functions in miRISC maturation or function. Moreover, by binding an essential repertoire of miRNA guides, the ALG-1(H751) protein, which is incapable of target repression, sequesters a large set of miRNAs that would otherwise associate with ALG-2 to function in the absence of ALG-1 (Figure 6A). We reason that such a sequestration effect can account for the majority of the antimorphic effect of H751L, where the mutant exhibits a stronger phenotype than the *null* mutant, without a major impact on miRNA levels. For other *alg-*1 NDD mutations that substantially affect the levels of more miRNAs, the overall phenotype could reflect a combination of both altered miRNA levels and sequestration of miRNAs in defective miRISC.

**Figure 6.**
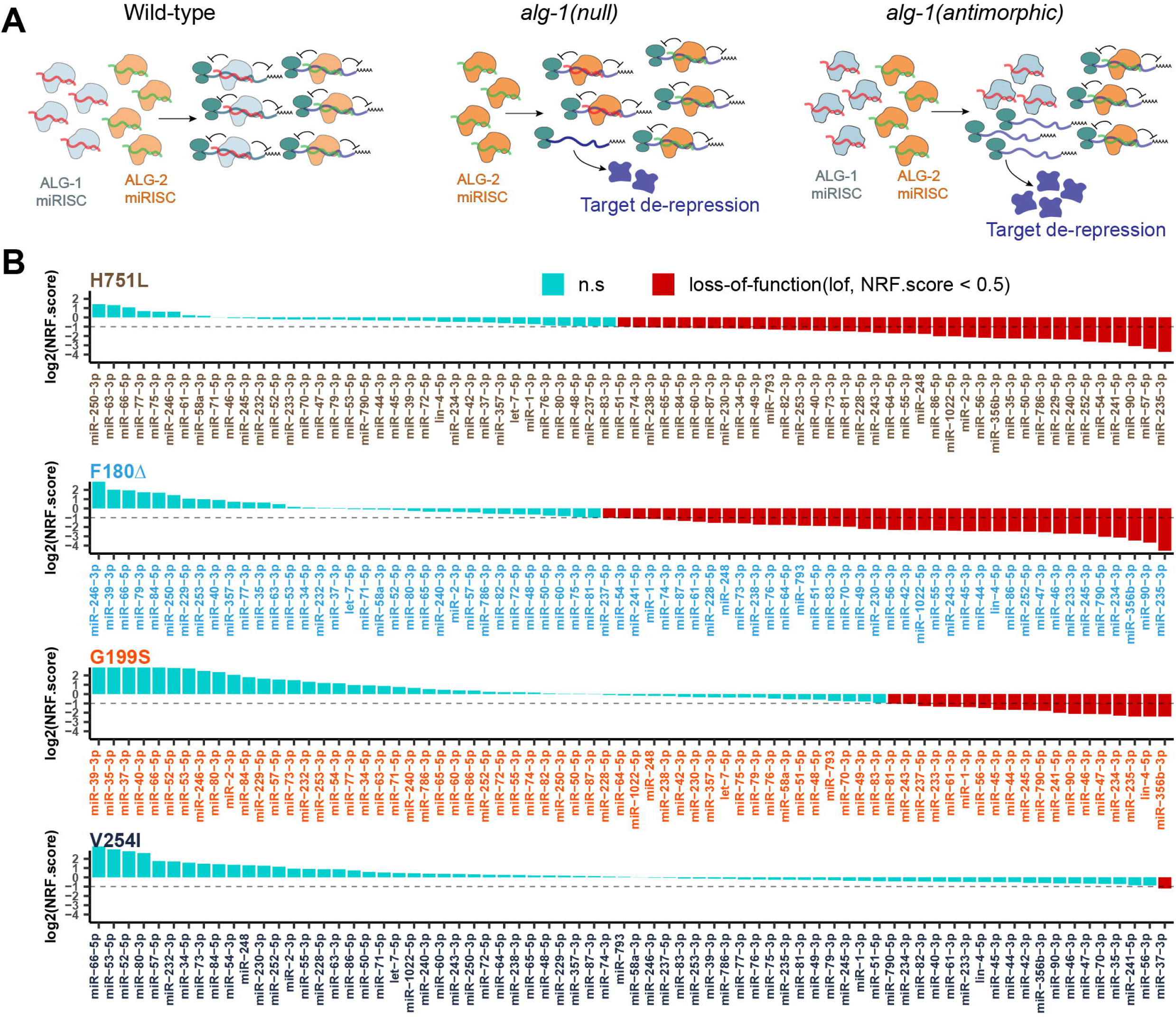
Antimorphic ALG-1(NDD) miRISC may sequester miRNAs into non-functional complexes, leading to a greater miRNA loss-of-function than in the absence of ALG-1. **A.** Illustrative models of WT and NDD ALG-1 miRISC activity, with ALG-1 NDD miRISC sequestering functional miRNAs away from the ALG-2 miRISC. Proposed miRISC activity is shown for *alg-1* WT, *alg-1(null),* and *alg-1(antimorphic)* genotypes. **B.** Putative <u>n</u>et repressive functionality score (NRF.score) of the most abundant miRNAs in the *alg-1* NDD mutants (min.RPM >15). Only the guide strands of miRNA were analyzed. miRNAs with NRF.score < 0.5 are defined as having a *lof* NRF.score. See also Table S4.

Following the model above, we propose that each of the *alg-1* NDD mutations cause a certain amount of net loss-of-function (*lof*) for specific miRNAs that is a combination of two effects: the reduction in the overall level of that miRNA and the sequestration of that miRNA into non-functional miRISC. To capture these two components in a single numerical estimate of *lof* for individual miRNAs in each *alg-1* NDD mutant, we derived a <u>n</u>et repressive functionality score (NRF.score; see Materials and Methods), which represents the proportional target repressive functionality of that particular miRNA in the mutant compared to the WT (NRF.score_WT_ = 1; NRF.score_null_ =0). The NRF.score captures the contribution from miRNA level reduction by incorporating input miRNA fold change compared to the WT. Meanwhile, the NRF.score also captures the contribution from sequestration of miRNAs in defective ALG-1 miRISC by incorporating two values: 1) the enrichment of the miRNA co-immunoprecipitation with ALG-1 in the mutant compared to the WT, and 2) the intrinsic function of the mutant ALG-1 protein determined by the penetrance of the lethality of *alg-1* mutant in *alg-2(null)* genetic background (Figure 1H).

We calculated the NRF.score of the 76 most abundant miRNAs (minimum RPM >15) and found that the numbers of miRNAs with NRF.score below an arbitrary threshold for *lof* (NRF.score < 0.5) were 44 for H751L, 45 for F180Δ, 22 for G199S, and 1 for V254I (Figure 6B, Table S4). Notably, by modeling overall miRNA functionality as sequestration in combination with perturbed levels, the NRF.score identified a larger number of miRNAs functionally affected by the H751L mutation, potentially reconciling the disconnection between the strong *alg-1(H751L)* phenotype compared to the mild effects of miRNA abundances.

To test whether modeling miRNA function according to NRF.score is consistent with the observed translatome disruption, we identified sets of putative disrupted miRNA targets for each mutant (145 for H751L, 396 for F180Δ, 81 for G199S, and none for V254I) that were translationally up-regulated and that also contain predicted target sites for the miRNAs with a *lof* NRF.score in that mutant. For the H751L, F180Δ, and G199S mutants, the corresponding putative disrupted miRNA targets were statistically enriched among all translationally up-regulated genes, compared to the target genes of just down-regulated miRNAs (Figure S5). These results support the model that the expression of mutant ALG-1 NDD protein in *C. elegans* can cause miRNA loss-of-function by the combined effects of disrupted miRNA biogenesis and sequestration of miRNAs in defective ALG-1 miRISC.

### The *alg-1* NDD mutations have distinct impacts on translational repression and mRNA abundance

miRNA-mediated post-transcriptional gene regulation can occur via translational repression and/or mRNA destabilization, such that impaired miRNA activity can manifest as increased translational efficiency (TE) and/or increased abundance of target mRNAs, respectively [33, 39]. To assess how the *alg-1* NDD mutations affect these two modes of target repression, we analyzed our ribosome profiling results in conjunction with RNA-seq analysis of total mRNA (Table S2). For the RNA-seq, we employed ribosomal RNA depletion for mRNA enrichment to ensure the quantitative recovery of all mRNAs regardless of poly(A) status [40]. The translational efficiency (TE) of each transcript was calculated by normalizing the RPF values with mRNA abundance [38]. We evaluated the TE and mRNA abundance of genes that were up-regulated in the translatome and that also contain predicted target sites for miRNAs with a *lof* NRF.score in each mutant. These genes were categorized into three de-repression modes: (a) genes that exhibit statistically significant up-regulation in TE but no significant change in mRNA abundance, referred to as “TE up”; (b) genes that exhibit statistically significant up-regulation in mRNA abundance but no significant change in TE, referred to as “mRNA up”; (c) genes that exhibit statistically significant up-regulation in both TE and mRNA abundance, referred to as “both up”(Figure 7B). We found that in the F180Δ, G199S, and H751L mutants, 79.2%, 79.1%, and 34.5% of the translationally up-regulated targets of *lof* miRNAs exhibit a statistically significant increase in TE and/or mRNA abundance (“TE up”, “mRNA up” or “both up”) (Figure 7C). Interestingly, 61.0% of these genes in the H751L mutant were de-repressed with increased translational efficiency without a significant change of mRNA abundance (“TE up”), whilst only 13.7% and 6.2% for the F180Δ and G199S mutants were de-repressed via “TE up” mode (Figure 7C).

**Figure 7.**
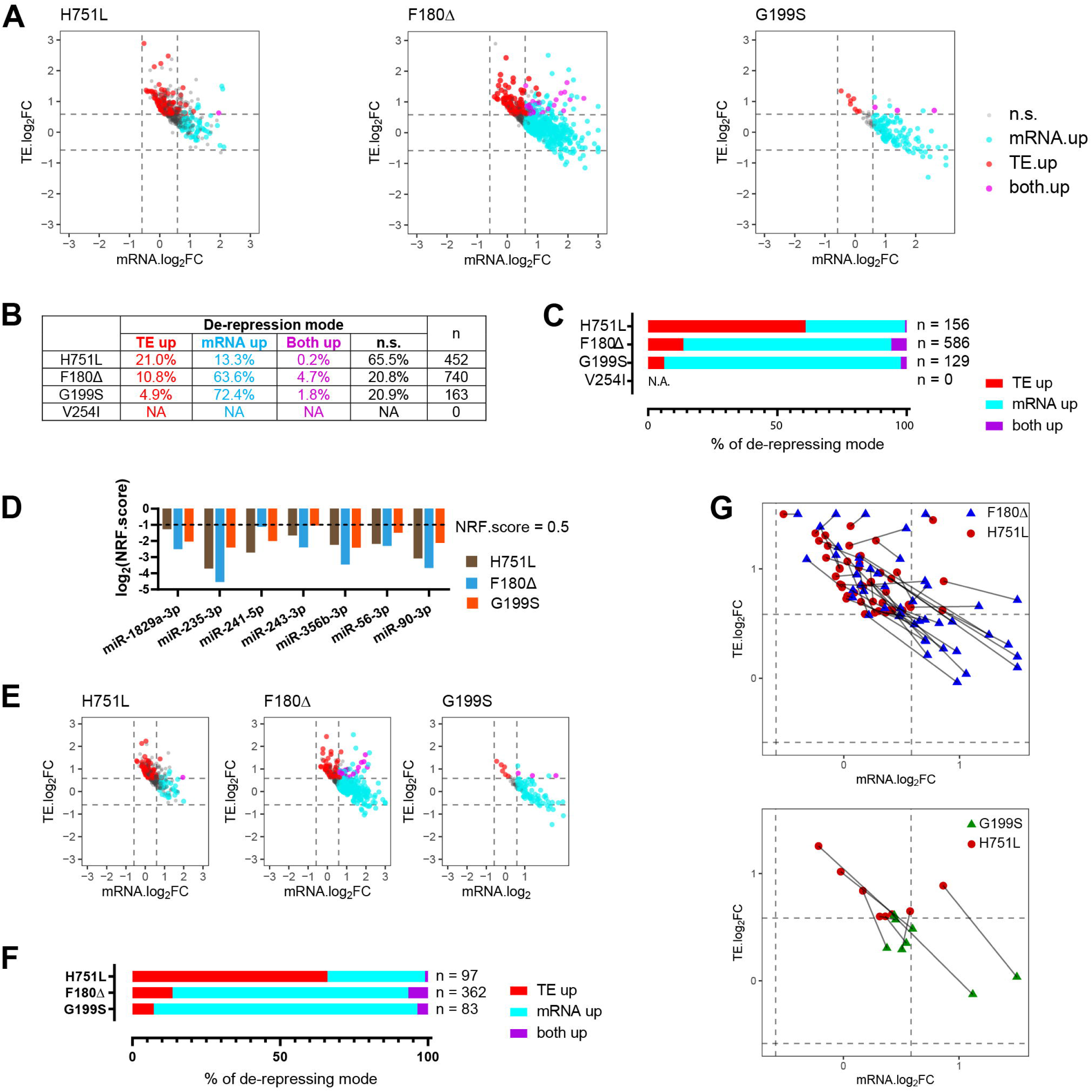
The *alg-1* NDD mutations have distinct impacts on gene target repressing modes based on translational efficiency and mRNA abundance. **A.** Fold changes of mRNA abundance and translational efficiency (TE) of genes that are significantly up-regulated and contain target sites of miRNAs with *lof* NRF.score (< 0.5). Cyan, genes with significantly increased mRNA abundance (|FC| > 1.5, p.adj < 0.1 by *DEseq2*). Red, genes with significantly increased TE (|FC| > 1.5, p < 0.1 by *Student t-test*). Magenta, genes with both significantly increased TE and mRNA abundance. **B.** Summary of the de-repression modes of genes that are translationally up-regulated and contain target sites for miRNAs with *lof* NRF.score. **C.** Distribution of the de-repression modes of the genes in (B) with significantly up-regulated TE and/or significantly up-regulated mRNA abundance. **D.** NRF.score of miRNAs that are down-regulated with statistical significance in both F180Δ and H751L mutants. **E.** Fold changes of mRNA abundance and TE of genes that are translationally up-regulated and contain target sites of miRNAs in (C). **F.** Distribution of the de-repression modes of the genes in (E) with significantly up-regulated TE and/or significantly up-regulated mRNA abundance **G.** Fold changes of mRNA abundance and TE of genes that contain targets sites of miRNAs in (C) and were simultaneously perturbed in both *alg-1(*F180Δ) and *alg-1(*H751L) or both *alg-1(*G199S) and *alg-1(*H751L) mutants.

The contrast in TE disruption bias associated with H751L could indicate that different ALG-1 NDD mutations can have selective disruptions of intrinsic functionalities of the ALG-1 protein, leading to distinct effects on the downstream target repression mechanism. However, an alternative explanation could be that the H751L mutation happens to preferentially disable the activity of a subset of miRNAs that are enriched for those with TE-regulated targets. To distinguish between these possibilities, we selected the set of miRNAs with a *lof* NRF.score in all F180Δ, G199S, and H751L mutants and analyzed the targets repression of these commonly affected miRNAs (Figure 7D). We found that the targets of the commonly affected miRNAs also showed a similar TE-only enrichment in the H751L mutant compared to F180Δ and G199S (Figure 7E-F).

Additionally, we found that the TE de-repression bias of H751L applies even to specific target genes. We analyzed the subset of 46 genes with “TE up” de-repression mode in the H751L mutant and compared the de-repression modes of these identical genes in the F180Δ or G199S mutants. We found 44 of the 46 genes were also de-repressed in F180Δ, and 25 out of those 44 genes exhibited a shift of the mode of de-repression from H751L to F180Δ. Similarly, 8 of the 46 genes de-repressed in H751L were also up-regulated in G199S, and all 8 genes exhibited a mode shift from H751L to G199S (Figure 7G). This suggests that the distinction in the target de-repression mode associated with individual NDD mutations is directly related to ALG-1 protein function, and not an indirect effect of selective disruption of unique sets of miRNAs or targets. We thus conclude that the H751L mutation may directly impair the target repression functionality of ALG-1 in a way distinct from the F180Δ and G199S mutations, perhaps reflecting discrete functions of the mutated amino acids and/or different roles in target repression mode for the PIWI and L1 domains where these mutations localize.

### The *alg-1* NDD mutations can perturb the expression of genes with human orthologs expressed in brain translatomes and/or related to NDD

The documentation of the hAGO1 mutations in human NDD patients raises the question of whether the perturbed genes in *C. elegans* NDD mutants include genes whose human homologs could be related to the pathogenesis of NDD. We examined the homology between the perturbed genes in *C. elegans alg-1* NDD mutants and the genes translationally expressed in human central nervous system [41, 42]. We found that among the *C. elegans* genes that were translationally perturbed in the *alg-1* NDD mutants, 262 genes for F180Δ, 61 genes for G199S, 6 genes for V254I, and 79 genes for H751L have human orthologs which are expressed in human brains translatomes (Figure 8A-B) [41, 42]. 55% for F180Δ, 34% for G199S, 0% for V254I, and 58% for H751L of these genes have target sites for the miRNAs with *lof* NRF.score (Figure S7). Of the neuronally-expressed human/worm homologous genes disrupted in *C. elegans alg-1* NDD mutants, 52 genes for F180 Δ, 13 genes for G199S, 3 genes for V254I, and 16 genes for H751L have been reported to be genetically associated with human NDDs with ID symptoms and definitive sysNDD entry [41, 43] (Figure 8C, Table S5). This observation suggests that the perturbation of these genes may contribute to the clinical manifestations observed in NDD patients from whom the mutations were identified.

**Figure 8.**
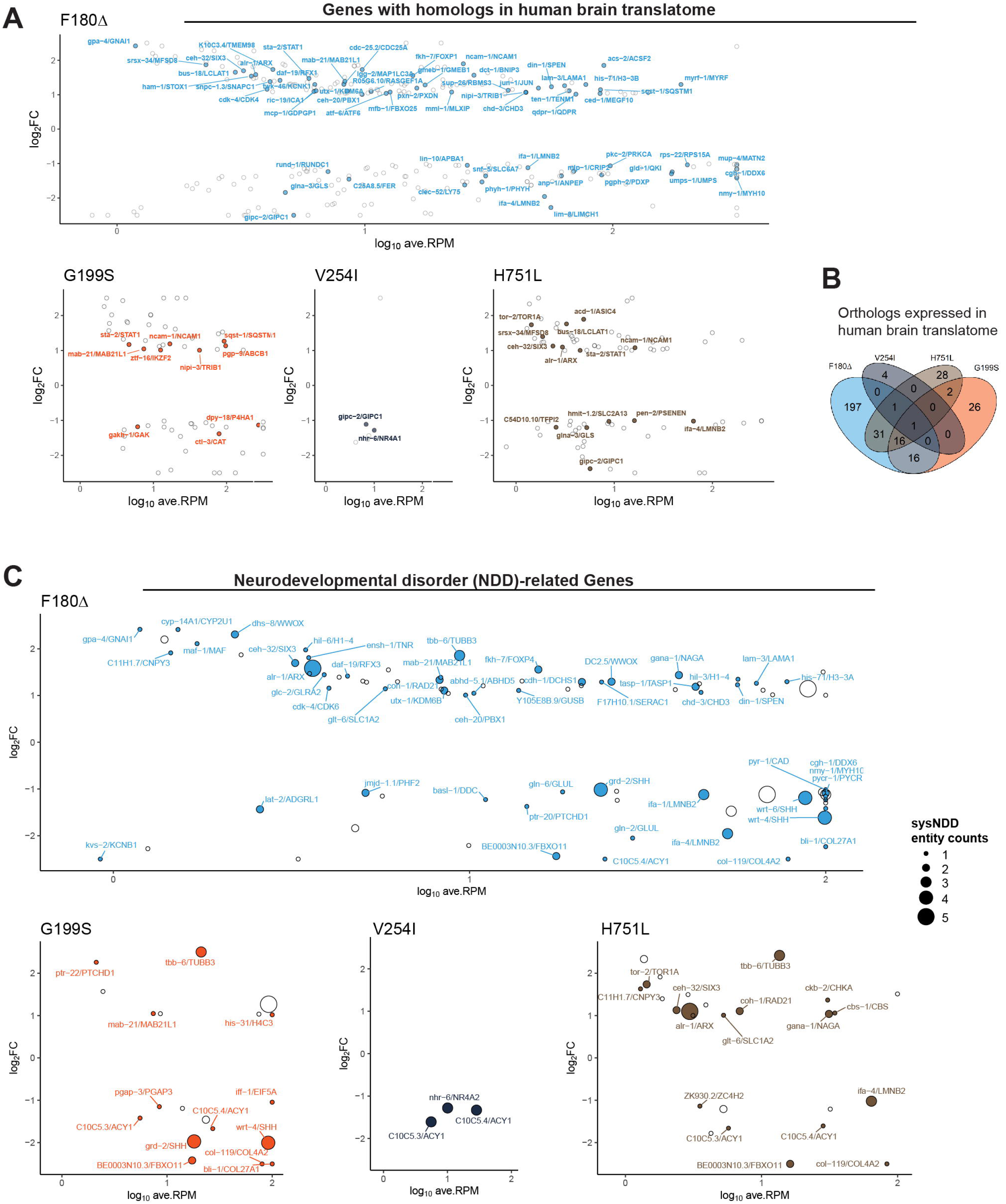
The *alg-1* NDD mutations can perturb genes with human orthologs expressed in the brain and human orthologs related to NDD. **A.** MA plots for the translational levels of up-regulated *C. elegans* genes which have human orthologs with brain translatome expression. Text-labeled and colored points indicate genes that are also expressed in the *C. elegans* nervous system [42]. The labels are formatted as *Cel_gene_symbol / Hsa_gene_symbol*. See also Table S5. **B.** Venn diagram for genes that are translationally up-regulated in *C.elegans* and have human orthologs expressed in brain translatome [42]. **C.** MA plots for translationally perturbed genes that have human homologs with sysNDD curation [43]. Solid and text labeled dots indicate genes that contain definitive sysNDD entity. Dot radius indicates the sysNDD entity counts. See also Table S5.

### The translatome perturbation in the *alg-1* NDD mutants may trigger stress-related responses due to proteome imbalance

Protein homeostasis (proteostasis) is tightly controlled and critical for normal cellular physiology. An imbalance in the proteome induced by genetic or other perturbations can impair proteostasis and contribute to pathogenesis [44, 45]. In normal cells, proteome imbalance elicits the activation of stress response pathways to restore proteostasis [46, 47]. For example, a recent study shows that aging-induced proteome imbalance in *C. elegans* can trigger stress responses due to abnormal protein aggregation [48]. In the *alg-1* NDD mutants, a large proportion of protein-coding genes have been translationally perturbed, especially in the F180Δ and H751L mutants (9.0% of total protein-coding genes for F180Δ and 3.4% for H751L). Accordingly, we found that stress-related genes are significantly enriched in the translationally up-regulated genes in all four NDD mutants, suggesting that the perturbation of translatomes may be leading to proteome imbalance, which consequently triggers stress responses (Figure 9A, S7A) [49, 50]. The F180Δ, G199S, and V254I mutants also exhibited up-regulation of small heat shock protein (HSP) genes, which encode chaperons that buffer insoluble protein aggregation [51] (Figure S7B), and the unfolded protein response (UPR)-related genes are also statistically enriched in the translationally up-regulated genes for all of the NDD mutants (Figure 9B), indicating that stress responses may be triggered by the misfolding and aggregation of proteins expressed at abnormally high levels in the mutants. Meanwhile, no significant changes were seen for the expression levels of large heat shock proteins orthologous to HSP70/HSP90, and no global up-regulation of proteosome components and proteolysis-related genes [52, 53] was observed (Figure S7C-D).

**Figure 9.**
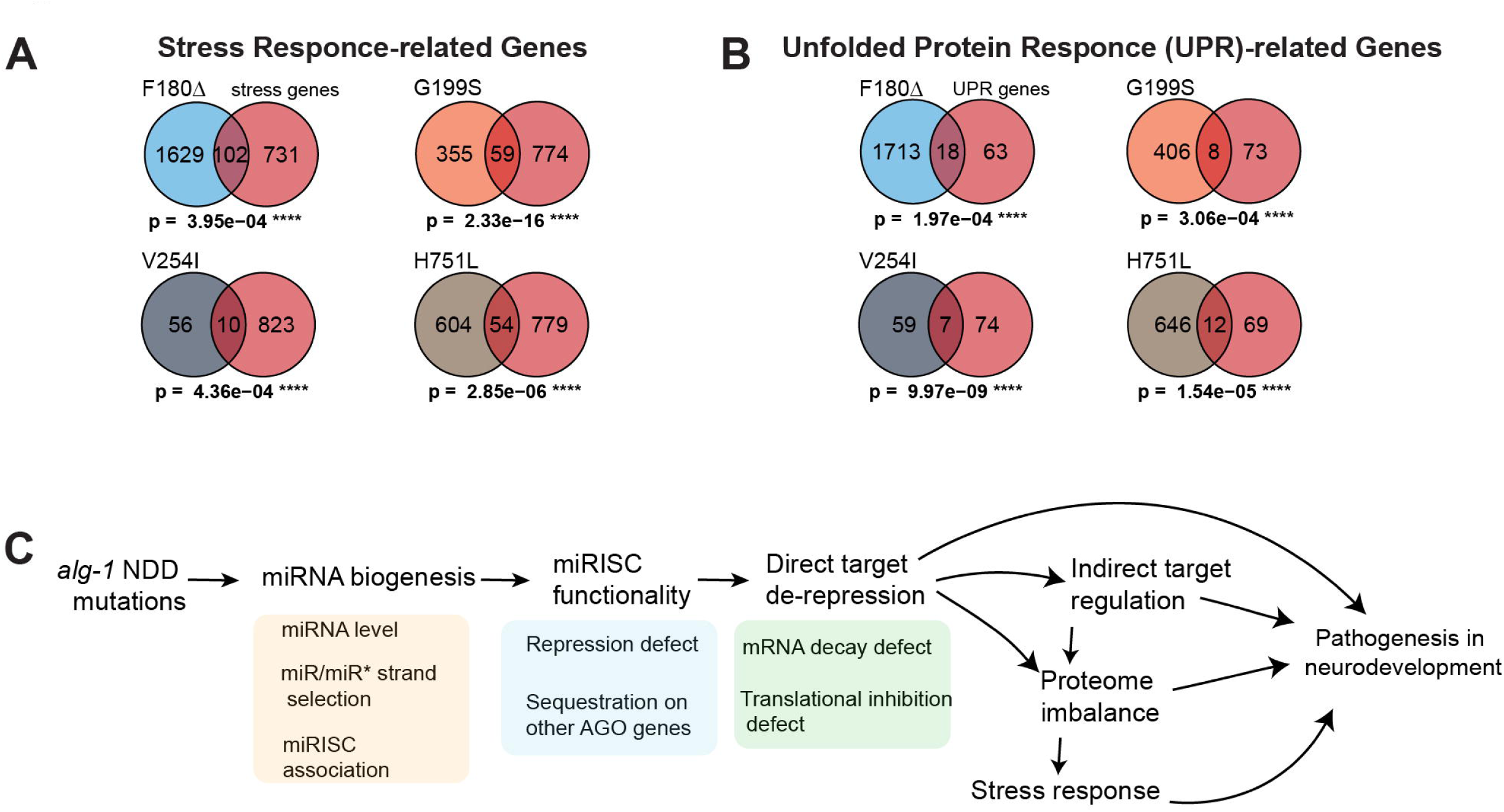
The *alg-1* NDD mutations can trigger stress response due to proteome imbalance. A-B. Hypergeometric tests for enrichment of stress-related genes (A) and unfolded protein response (UPR) (B) in translationally up-regulated genes in the *alg-1* NDD mutants [49, 50]. **C.** Summary of the possible contribution of the *alg-1*/*hAGO1* NDD mutations to the pathogenesis of NDD.

It is also noteworthy that the perturbation of small HSP genes was only observed for the F180Δ, G199S, and V254I mutants, but curiously, not in the H751L or *null* mutants (Figure S7B). The observation that the *alg-1* H751L and *null* mutant do not display the up-regulation of small HSP may suggest that proteome stress can be allelic specific and is not necessarily an inevitable consequence of disrupting miRISC function, but rather related to the particular repertoire of proteins dysregulated in the particular mutant.

## Discussion

### Modeling *hAGO1 de novo* coding variants in *C. elegans* enables the characterization of novel allele-specific Argonaute functions

Argonaute proteins of the Ago class contribute to miRNA biogenesis, as well as mRNA target recognition and repression [54]. Accordingly, depletion of AGO genes by RNAi or mutation can impair miRNA biogenesis and trigger miRNA target de-repression [6, 7, 23]. Structural studies of mammalian Argonautes and analyses of the evolutionary conservation of Argonaute amino acid sequences provide insights into the functional architecture of Argonautes and the functional importance of specific amino acids [6, 30, 32, 54, 55]. For example, D597, D669, and H807 are confirmed as key residues for the catalytic activity of slicing by hAGO2, and amino acids M47, D95, and F181 are known to critically contribute to the unwinding of miRNA duplex during the process of loading [56, 57]. Other functionally key residues have been revealed by forward genetic screens, exemplified by G553 and S895, for which mutations at the corresponding amino acids in *C. elegans* ALG-1 can impair miRNA biogenesis and guide-passenger strand selection [12, 29].

The recently described *de novo* mutations in *hAGO1* and *hAGO2* carried by NDD patients point to the significance of the corresponding amino acids in AGO function [20, 21]. It is noteworthy that most of the amino acids mutated in these patients, although phylogenetically conserved, had not been explicitly linked to AGO functions by previous studies. Thus, genetically modeling specific AGO mutations identified in human patients in an experimental animal promises to help elucidate novel functions of AGO proteins.

In this study, we chose four *hAGO1* mutations to model in *C. elegans* ALG-1 based on either of two criteria: (1) they were documented in multiple independent families (F180Δ, G199S, V254I) or (2) the mutation is adjacent to a previously identified phenocritical residue (H751L). We find that the *alg-1* NDD mutant proteins are expressed in *C. elegans,* associate with miRNAs, and cause visible phenotypes stronger than *alg-1 null*. Further, we find that the *alg-1* NDD mutations disrupt miRNA profiles in *C. elegans*, and cause patterns of translatome perturbation distinct from *alg-1 null* mutant. These properties are consistent with the antimorphic mutation model applied to a previously-described class of *C. elegans alg-1* mutations [12].

Notably, the severity of the *C. elegans* developmental phenotypes of the *alg-1* NDD mutants is consistent with symptom severity in the NDD patients. In *C. elegans,* F180Δ and H751L strongly impaired the viability, vulval integrity, and larval-to-adult differentiation of hypodermal cells, while the G199S mutation conferred more moderate phenotypes and the V254I exhibited nearly undetectable phenotypes. Similarly, the H751L patients (monozygotic twins, n = 1) exhibit severe ID with growth delay, microcephaly, speech impairment, motor delay, feeding difficulty, facial dysmorphia, and F180Δ patients (n = 9) exhibit mild-to-severe ID and motor delay, and some patients developed epilepsy, facial dysmorphia, growth retardation [20]. In contrast, the G199S patients (n = 9) exhibit mild-to-moderate ID with speech impairment, epilepsy, motor delay, and facial dysmorphia for some patients. Meanwhile, the V254I patients (n = 2) exhibit the least severe symptoms with mild ID, speech delay, epilepsy, and hyperactivity but no motor delay or additional features (no growth retardation or MRI anomalies) [20]. This broad correspondence of phenotypic severity between the two systems suggests that the mutations may impair the Argonaute protein in a similar fashion, highlighting the utility of model organisms to study human genetic disorders.

Interestingly, the four NDD mutations displayed distinctive impairment of AGO functions in *C. elegans.* Additional to the distinctive developmental phenotypes discussed above, these mutations also differed in their effects on miRNA biogenesis and gene expression in several ways: (1) The severity of perturbations in miRNA levels and translatomes varied between different mutants; (2) The profiles of disturbed miRNAs and gene expression were distinct for each mutant; (3) The mutations displayed allele-specificity in their relative impacts on target mRNA abundance versus translational efficiency. Furthermore, the severity of phenotypes, miRNA perturbations, and translatome perturbations was not strictly correlated among the four *alg-1* NDD mutations. The distinctions in developmental and molecular phenotypes between different NDD mutations in *C. elegans* suggest the mutated amino acids have differing mechanistic impacts on *in vivo* functions of ALG-1 protein.

Currently, we do not have direct structural or biochemical evidence to illuminate specifically how the different NDD mutations modeled here could impair AGO protein functions. However, our results suggest possible mechanisms in light of current structural models. According to the current understanding of hAGO2 crystal structure, H751 resides inside the PIWI-MID channel where the miRNA seed region duplexes with the target RNA. The imidazole group in the H751 side chain is close to the backbone of g5 and g6 nucleotides of the miRNA and likely contacts the backbone phosphates via hydrogen bonds and electrostatic forces [32]. In H751L mutant ALG-1, the change from histidine to leucine alters the charge and hydrophobicity of the side chain and thereby may impair the interaction between the residue and the miRNA. Interestingly, we found that mutations at the two residues preceding H751 (S750F, and C749Y) can also strongly impair ALG-1 function, as revealed by the strong developmental phenotypes of the mutants. Since each of the S750F and C749Y mutations alters the hydrophobicity and spatial size of the side chains, these mutations may change the positioning of H751 and consequently prohibit it from interacting properly with the miRNA. The changes in the side chain size and hydrophobicity in these mutations may also result in allosteric distortion of ALG-1 protein and impair function associated with more distant domains of ALG-1.

A recent study of *Arabidopsis thaliana* Argonaute AtAGO10 suggests that a ß-hairpin of L1 domain, which is conserved in eukaryotic AGOs, contacts the t9-t13 of target RNA by electrostatic forces and consequently coordinates the pairing between 3’ non-seed region of miRNA and target [58]. Interestingly, the L1 ß-hairpin includes the residue homologous to *hAGO1* F180 and is sterically adjacent to the residue homologous to *hAGO1* G199 (Figure S8). Moreover, the structure of the human AGO2::miRNA::target complex suggests that the ß-hairpin resides sterically adjacent to t11-t13 of the target RNA (Figure S8B) [31]. Recent genetic and biochemical studies have shown that such 3’ pairing, especially at t11-t13, can be critical to the proper regulation of certain miRNA/targets [38, 59]. Thus, although F180 and G199 residues do not directly contact the 3’ duplex of miRNA/target, the F180Δ and G199S mutation may disrupt 3’ pairing by distorting the sub-regional conformation of the L1 ß-hairpin or hinder its movement, and consequently impair target repression, especially for the miRNA/target interactions that requires 3’ pairing. Moreover, extrapolating from the AtAGO10 structure in the slicing configuration, where the L1 ß-hairpin interacts with the non-guide strand of the helical miRNA::target duplex, it is possible that the F180Δ and G199S mutations could disrupt interactions of the L1 ß-hairpin with the passenger strand of AGO::pre-miRNA complexes. Such a hypothetical interaction is consistent with our results that the F180Δ and G199S mutations disrupt of miRNA biogenesis.

In contrast, the V254 residue is neither directly interacting with the miRNA::target duplex nor involved in sub-regions with specific functions that have been structurally or biochemically characterized. According to the current structure, the V254 side chain is exposed on the surface of the AGO protein, enabling it to potentially contact other protein factors. Thus, the V254I mutation may impair AGO protein function by impacting inter-molecular interactions with other proteins.

### Molecular mechanisms of the *antimorphic* effect of *alg-1* NDD mutations

It is striking that the reported *hAGO1* and *hAGO2* NDD mutations are mostly single amino acid changes [20-22]. The rarity of frameshift, truncation, or large deletions suggests that single-gene *hAGO null* mutations are either not tolerated or do not cause observable symptoms. For *hAGO2*, the rarity of *de novo null* mutations in NDD patients could be due to the critical contribution of *hAGO2* to miR-451 biogenesis, which is essential for erythropoiesis and erythroid homeostasis in mammals [23, 60, 61]. Interestingly, *null* mutations of *hAGO1* in NDD patients have been reported as large deletions that also delete the nearby *hAGO3* and *hAGO4* genes [18, 19]. This suggests that *hAGO1*, *hAGO3*, and *hAGO4* are redundant, and that diagnosable phenotypes do not arise unless the activity of multiple AGO genes is defective. If this is so, how could the reported point mutations in *hAGO1* or *hAGO2* result in phenotypes? This question motivates the antimorphic model for the action of the *hAGO1* NDD mutations, wherein the mutant AGO protein impairs miRNA activity by competing with otherwise redundant paralogous AGO proteins. In this model, the mutant AGO protein is expressed and can associate with miRNAs but is functionally defective in target repression. Consequently, a large fraction of miRNAs in the cell is sequestered in non-functional complexes, depleting the supply of miRNAs available to paralogous AGO proteins.

In this study, we show that the H751L and F180Δ mutations are antimorphic for *alg-1* in *C. elegans* because the mutants exhibit heterochronic phenotypes with greater penetrance than that of the *alg-1 null* mutant. The antimorphic effect of H751L and F180Δ is similar to the results from Zinovyeva et al. 2014 where the *alg-1(ma192)* (corresponding to S750F for *hAGO1*) and *alg-1(ma202)* (corresponding to G571R for *hAGO1*) mutations exhibit a similar antimorphic effect on *C. elegans alg-1* [12]. These parallels suggest that these human and *C. elegans* AGO mutations may similarly result in the expression of mutant Argonaute proteins that can associate with miRNAs but that are functionally defective in target repression, and hence sequester miRNAs in non-functional miRISC.

The sequestration model is consistent with our results. In particular, we found that the H751L mutation only mildly disturbed the profiles of total miRNA expression or the profiles of miRNA co-immunoprecipitated with mutant ALG-1. The mild disturbance of miRNA profiles suggests that ALG-1^H751L^ supports essentially normal miRNA biogenesis and miRISC assembly, but that the ALG-1 miRISC may be defective in target recognition and repression. Considering the possibility that the H751L mutation could impair the interaction of ALG-1 with the backbone of the miRNA seed region (Fig. 2A), it is possible that the H751L mutation prevents the functional interaction of miRISC with targets. Thus, in the case of H751L, the antimorphic effect could result largely from the sequestration of miRNAs in miRISC complexes that are unable to bind targets. We also note that, in principle, antimorphic mutations could disrupt miRISC function at step(s) after target binding. In this scenario, non-functional miRISC would bind the targets and competitively inhibit access to the target by miRISC containing other AGO proteins, essentially exerting a blocking effect in addition to the sequestration effect.

Furthermore, *alg-1* antimorphic mutants can exhibit potent disruption of guide/passenger strand selection [29], which could in principle result in essentially neomorphic miRNA phenotypes in cases where the normally degraded passenger strand accumulates to functional levels. If the mutant ALG-1 miRISC were to retain partial function -- as is the case for the G199S and V254I mutants which are viable in *alg-2 null* background -- then ALG-1 miRISC containing hyper-abundant passenger strands could repress target genes which are not supposed to be regulated by miRNAs. Such neomorphic effects can also contribute to the production of phenotypes distinct from the *null* mutant.

### The pleiotropy of AGO functions and the pathology of the AGO NDD mutations

In this study, we show that the *alg-1* NDD mutations can globally perturb translatome in *C. elegans*. Although the severity of translatome perturbation varied among different mutations, the stronger mutations were observed to disrupt remarkable proportions of the translatome. Meanwhile, some NDD mutants also exhibit strong developmental defects, including F180Δ and H751L, which are embryonically arrested in *alg-2 null* genetic background. These global perturbations in gene expression and severe developmental defects are consistent with the extensive pleiotropy of AGO-mediated miRNA regulation.

The perturbation of gene expression caused by *alg-1* NDD mutations was not only extensive but also remarkably distinct, with each mutant exhibiting sets of disrupted genes that were unaffected in the other mutants. Extending these observations to hypothetical effects of the corresponding mutations in *hAGO1*, it is reasonable to suggest that similar extensive and partially allele-specific gene expression disruptions could occur in the patients with *hAGO1* or *hAGO2* NDD mutations. Here, a key question arises: could NDD pathology result from the dis-regulation of a small set of specific genes whose over-expression is causative for NDD and that happen to be dis-regulated in common by all the mutations? We suggest that this scenario is possible and warrants further investigation in mammalian systems. However, the distinctions among the *C. elegans alg-1* NDD mutations, both in distinct repertoires of disrupted miRNAs and distinct profiles of downstream gene perturbations, suggest that NDD pathology in human patients could reflect, at least in part, the emergent physiological and developmental consequences of broad dis-regulation of gene expression networks. This supposition is consistent with the observation that even among patients carrying identical *hAGO1* or *hAGO2* NDD mutations, there is still variability of clinical manifestation compared to some other NDD syndromes [20, 21, 62].

The AGO NDD mutations can be thought of as triggering cascades of gene expression dis-regulation (Figure 9C). We found that the *alg-1* NDD mutations can disrupt the processing, loading, and/or function of multiple miRNAs. Since individual miRNAs can have dozens to hundreds of targets, it is expected that the immediate impact of *alg-1* NDD mutations would include de-repression (or neomorphic repression in the case of altered miRNA strand selection) of a large set of direct targets. Perturbations of direct miRNA targets, particularly regulatory gene products such as RNA binding proteins and transcription factors, should in turn lead to amplified downstream disruptions of gene regulatory networks. The disrupted gene sets, including direct miRNA targets and the indirectly affected downstream genes, can include sets of genes expressed in the nervous system and/or with human homologs genetically linked to NDD-related phenomena (Figure 8).

Among the potential physiological impacts of global dis-regulation of gene expression in NDD mutants, it seems appropriate to consider the cellular and organismal stress of proteome imbalance. We observed a statistically enriched up-regulation of the expression of stress-related genes in some of the *C. elegans alg-1* NDD mutants. This up-regulation includes small heat shock proteins, which is indicative of a proteome imbalance-induced protein aggregation [48, 52, 63]. Thus, one physiological trigger underlying the pathological effects of the AGO NDD mutations could originate from cellular and organismal responses to the disturbance of proteostasis caused by global perturbation of gene expression (Figure 9C).

## ACKNOWLEDGMENT

We thank the members of Ambros Lab, Mello Lab, and Zinovyeva lab for the project discussion. We thank Brittany Morgan and Francesca Massi for commenting on the structural modeling. This research was supported by funding from NIH grants R01GM088365, R01GM034028, R35GM131741 (VA), and R35GM124828 (AZ). Some *C. elegans* strains were provided by the CGC, which is funded by the NIH Office of Research Infrastructure Programs (P40 OD010440).

## Author Contributions

Conceptualization: YD, AP, AZ, VA; Methodology: YD, LL, GPP, AZ, VA; Formal analysis: YD, LL, GPP; Investigation: YD, LL, GPP; Resources: AZ, VA; Data curation: AZ, VA; Writing -original draft: YD, AZ; Writing -review & editing: YD, LL, AZ, AP, VA; Supervision: AZ, VA; Project administration: VA; Funding acquisition: AZ, VA.

## Declaration of Interests

The authors declare no competing interests.

## METHODS

### *C. elegans* culturing and synchronization

*C. elegans* were cultured on nematode growth medium (NGM) and fed with *E. coli* HB101. To obtain populations of synchronized developing worms, gravid adults were collected and washed twice with water. Pellets of centrifuged worms were treated with 5 ml 1N NaOH and 1% (v/v) sodium hypochlorite for 4 min with shaking to obtain embryos, and the embryos were rinsed with M9 buffer three times. The embryos were hatched in 10 ml M9 buffer at 20°C for 16-18 hrs with mild shaking. Hatched L1 larvae were transferred to plates at 30-50 worms per plate and replicate plates were cultured at 15°C, 20°C, or 25 °C for defined periods of time; samples of the population were examined by microscopy to confirm the developmental stages at the time of harvest.

### CRISPR/Cas9 targeted mutagenesis at the *alg-1* genomic locus

Templates for ssDNA HR donors with 45-60 nt flanking the mutated nucleotide(s) were obtained from IDT. To generate the V254 and H751L mutants, CRISPR/Cas9 RNP mixtures were injected into N2 animals at the following final concentrations: Alt-R Cas9 (1.9 µM, IDT, cat# 1081058), AltR_Cas-9_crRNA_dpy-10_cn64 (0.4 µM, IDT) [64], two AltR_Cas-9_crRNA_alg-1_H751/V254 crRNAs specific for the edited regions (0.6 µM each, IDT), Alt-R tracrRNA (1.6 µM, IDT, cat# 1072532), and *alg-1* H751L or V254I donor (160 ng/µL) in Cas9 RNP annealing buffer (1x, IDT, cat# 11010301). The Cas9 RNP mixtures were incubated at 37°C for 5 minutes, and spun down for 2 minutes at 14000 RCF prior to injections. To generate the F180Δ/G199S/C749Y mutants, CRISPR/Cas9 RNP mixtures were injected EG9615 animals which express transgenic Cas9 from *oxIs1091* integrated transgene [65] at the following final concentrations: AltR_Cas-9_crRNA_dpy-10_cn64 (0.86 µM, IDT), AltR_Cas-9_crRNA_alg-1_F180/G199/C749 crRNA specific for the edited regions (2.6 µM, IDT), Alt-R tracrRNA (3.5 µM, IDT, cat# 1072532), and 120 ng/µl ssDNA donor in 1X duplex buffer (Table S6).

F1 dumpy and/or non-dumpy animals were isolated from dumpy jackpot plates and genotyped by PCR and restriction digestion using HpyCH4IV (F180Δ, NEB R0619), HinfI (G199S, NEB R155S), AflIII (V254I, NEB R0541S), RsaI (C749Y, NEB N0167S) and DdeI (H751L, NEB R0175), followed by Sanger sequencing. Mutants were backcrossed with N2 at least twice to remove *dpy-10, oxIs1091,* or other potential background mutations.

### Phenotypic assays for vulva defects

The adult lethality which results from the rupture of the young adult animal at the vulva (burst) or matricide by offspring hatching in the uterus (bag) was scored after approximately 36 hrs (15°C), 24 hrs (20°C) or 16 hrs (25°C) of development (when at least 95% of the population had reached the adult stage). To score viable progeny per adult, young adults were transferred to a fresh plate every 12 hrs until those capable of laying eggs had completed egg-laying. Only hatched eggs were counted.

### Microscopy and heterochronic phenotypes

Differential interference contrast (DIC) and fluorescent microscopy were performed on Zeiss.Z1 or Leica DM6 B compound microscopes equipped with epifluorescence capabilities. *col-19::gfp* patterns were scored by 10X or 63X objective. Fluorescent images were obtained on Zeiss.Z1 equipped with ZEISS Axiocam 503 camera and processed by ImageJ FIJI [66].

### Total RNA preparation

Harvested worms were washed with M9 medium, centrifuged, and the worm pellets were flash-frozen in liquid nitrogen. The worm pellets were thawed and lysed by adding 4X volumes of QIAzol (Qiagen, Cat: 79306) and shaking vigorously at room temperature for 15 min. The total RNA was extracted by the addition of 0.85X volume chloroform, centrifugation, and recovery of the aqueous phase, which was then re-extracted with 1 volume phenol:chloroform:isoamyl alcohol (25:24:1, pH = 6.3). Total RNA was then precipitated by adding 1 volume of isopropanol and 1 µl GlycoBlue (Invitrogen, Cat: AM9516), followed by incubation at −80°C for at least 30 min, and recovery by centrifugation at 25,000 rcf for 10 min at 4°C. The supernatants were discarded, and the RNA pellets were subsequently washed twice with 70% (v/v) ethanol, air-dried at room temperature for 5 min, dissolved in RNase-free water, and stored at −80°C.

### ALG-1 immunoprecipitation

The synchronized fourth larval stage (L4) animals were collected and the worm pellets were flash-frozen in liquid nitrogen and stored at −80C until total protein lysate preparations. Protein lysates were obtained as previously described [67]. ALG-1 immunoprecipitation and Western blotting were performed as previously described [68].

### Small RNA sequencing

Total (“input”) lysates and “IP” samples were subjected to RNA preparation as described above. Purified RNA was subjected to gel-based size selection as previously described [69]. NEXTflex Small RNA Library Prep kit v3 (PerkinElmer, cat# NOVA-5132) was used to prepare libraries according to the manufacturer’s instructions, followed by size selection of final PCR products as previously described[69]. Libraries were sequenced using the Illumina Nextseq500 platform at the Kansas State University Integrated Genomics Facility.

Small RNAseq reads were checked for quality before and after filtering using FastQC v0.11.8 (https://www.bioinformatics.babraham.ac.uk/projects/fastqc). Cutadapt tool was used to clip the adapter sequence from 3’ end (-a ATCTCGTATGCCGTCTTCTGCTTG −e 0.1). Reads were split into libraries using fastx barcode splitter utility (http://hannonlab.cshl.edu/fastx_toolkit/index.html) and the remaining 3’ end and 5’ adapter sequences were clipped. The randomers were trimmed and the reads with a final length range of 17-29 nt were selected for further analysis. Reads were mapped to C. elegans genome (WS279) using bowtie v1.2.2 [70, 71] allowing three mismatches in the alignment. Mature miRNA expression was quantified using the miRDeep2 pipeline [72]. The DESeq2 package in R was used to perform differential expression analysis [73].

### miRNA site prediction

miRNA targets were predicted against the transcriptomic 3’ UTR sequences using TargetScanWorm version 6.1 [74]. Sites with full complementarity to g2-g7 of the miRNA seed were retained. Sites with a seed mismatch at g5-g8 accompanied by a full pairing of g13 through g16 were also retained. Differential expression analysis was performed by DESeq2, significance indicates FDR (<0.05).

### Ribosome profiling

Synchronized populations of developing worms were cultured at 20 °C for 45 hrs after feeding. Worms harvest, monosome preparation, ribosome protected footprint (RPF) cloning and data analysis were performed as previously described [38] except that the RPF libraries were prepared using NEBNext Multiplex Small RNA Library Prep Set for Illumina (NEB E7300). The trimmed RPF reads were mapped to *C. elegans* genome WS279 [75]. Genes with |FC| >2 and p.adj <0.1 (*DESeq2*) were considered translationally perturbed genes with statistical significance.

### RNA-seq and translational efficiency (TE)

Worm samples for RNA-seq were aliquoted from the ribosome profiling harvests before the lysis step and frozen separately. The mRNA was enriched by ribosomal RNA (rRNA) depletion as described in [40], with additional ASO oligos to deplete small recognition signal RNA (srpR). Library preparation and RNAseq data analysis were performed as described in [38]. To calculate the TE, a pseudo count of 0.1 were added to each gene for all samples. Genes with |FC| >1.5 and p.adj <0.1 (*DESeq2* for RNA abundance and Student’s t-test for TE) were considered as significantly perturbed genes. The ASO sequences can be provided upon request.

### Calculation of Net repressive functionality score (NRF.score)

The relative functionality of a given miRNA (miR*i*) in a particular *alg-1* mutant (mut.) is calculated as:

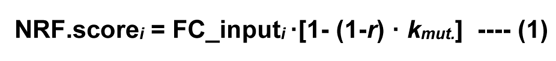

In equation (1), **FC_input*i*** indicates the fold change of total miR-*i* in the mutant, which is expressed as

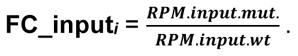

***r*** (0 ≤ *r ≤* 1) is the relative intrinsic functionality of mutant ALG-1 and is calculated by 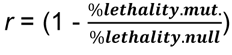 in the *alg-2* null genetic background (i.e., since the lethal phenotype of *alg-1(V254I);alg-2(0)* has a penetrance of 9.98%, *r*V254I is 0.9002). Thus **(1-*r*)** represents the proportion of inactivated repressing functionality of ALG-1.

***k*** (0 ≤ *k ≤* 1) is the enrichment of the miR-*i* that is co-immunoprecipitated with mutant *ALG-1*, which is expressed as 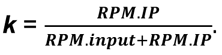. Thus, equation (1) can also be expressed as:

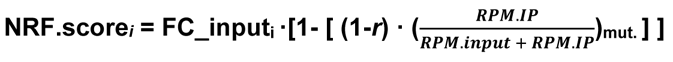

For the *alg-1* null mutant, r=0, therefore,

**NRF.score_null_ = FC_input_null_**

### Quantification and statistical analysis

p-values representation is as follow: 0.05-0.01(*); 0.01-0.001(**); 0.001-0.0001(***); <0.0001(****). The brood size/numbers of progeny phenotypes were analyzed by Student’s t-test (two-tailed, unpaired). The lethality and *col-19::gfp* expression phenotypes were analyzed by Fisher’s test. Error bars indicate mean ± SD. Significance tests were conducted with Prism 9.

## SUPPLEMENTAL FIGURE LEGENDS

**Figure S1.**
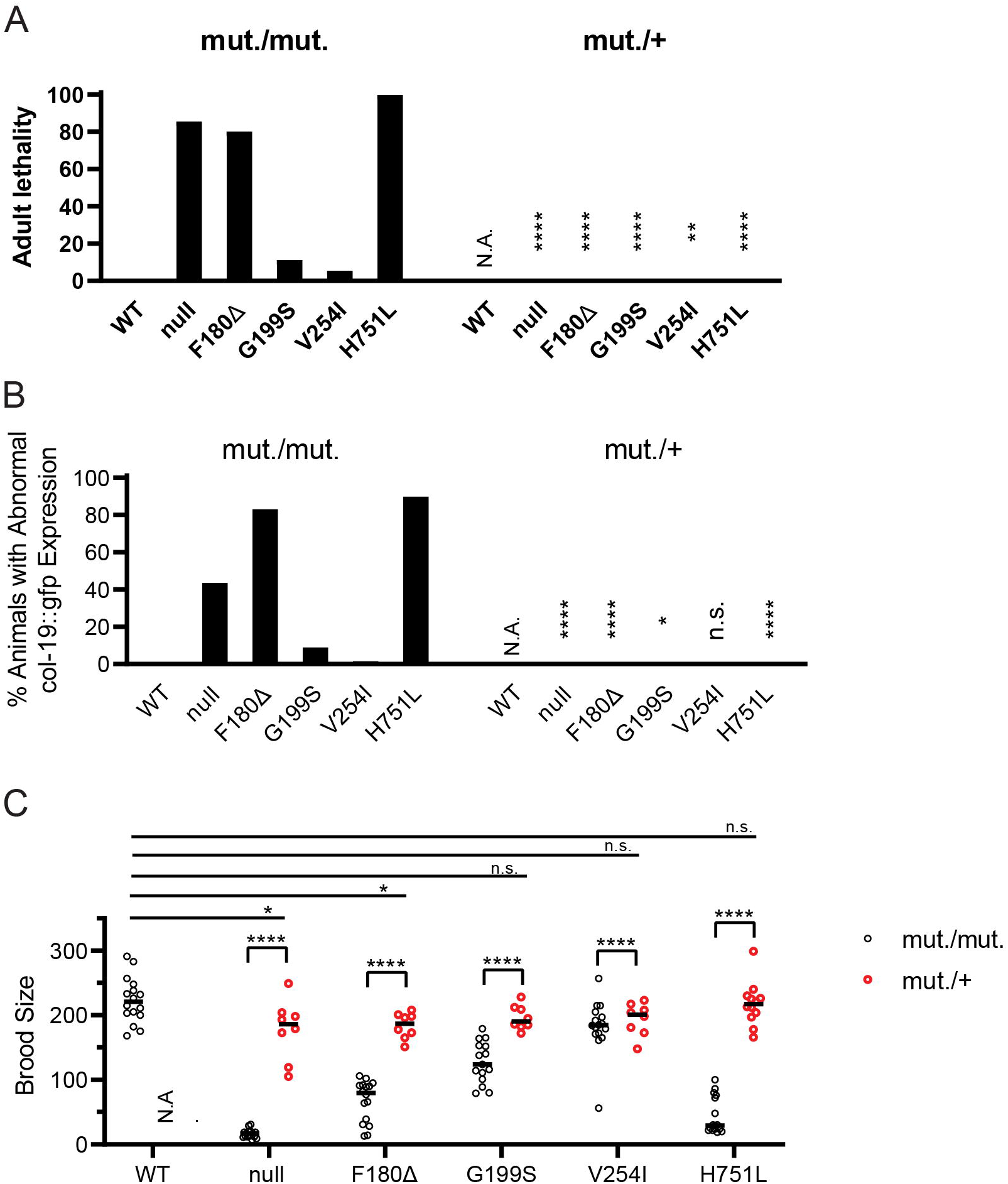
Effects of homozygous and heterozygous *alg-1* NDD mutations on vulval integrity and seam cell differentiation. **A-C.** The adult lethality (A), abnormal *col-19::gfp* (B), and numbers of progeny (C) phenotypes of the homozygous (mut/mut) and heterozygous (mut/+) *alg-1* NDD mutants. Phenotypes are scored at 25 °C. The statistical significance of lethality and abnormal *col-19::gfp* expression is analyzed by Fisher’s test. The statistical significance of brood size is analyzed by Student t-test (see Method). ****p≤0.0001, ***p≤0.001.

**Figure S2.**
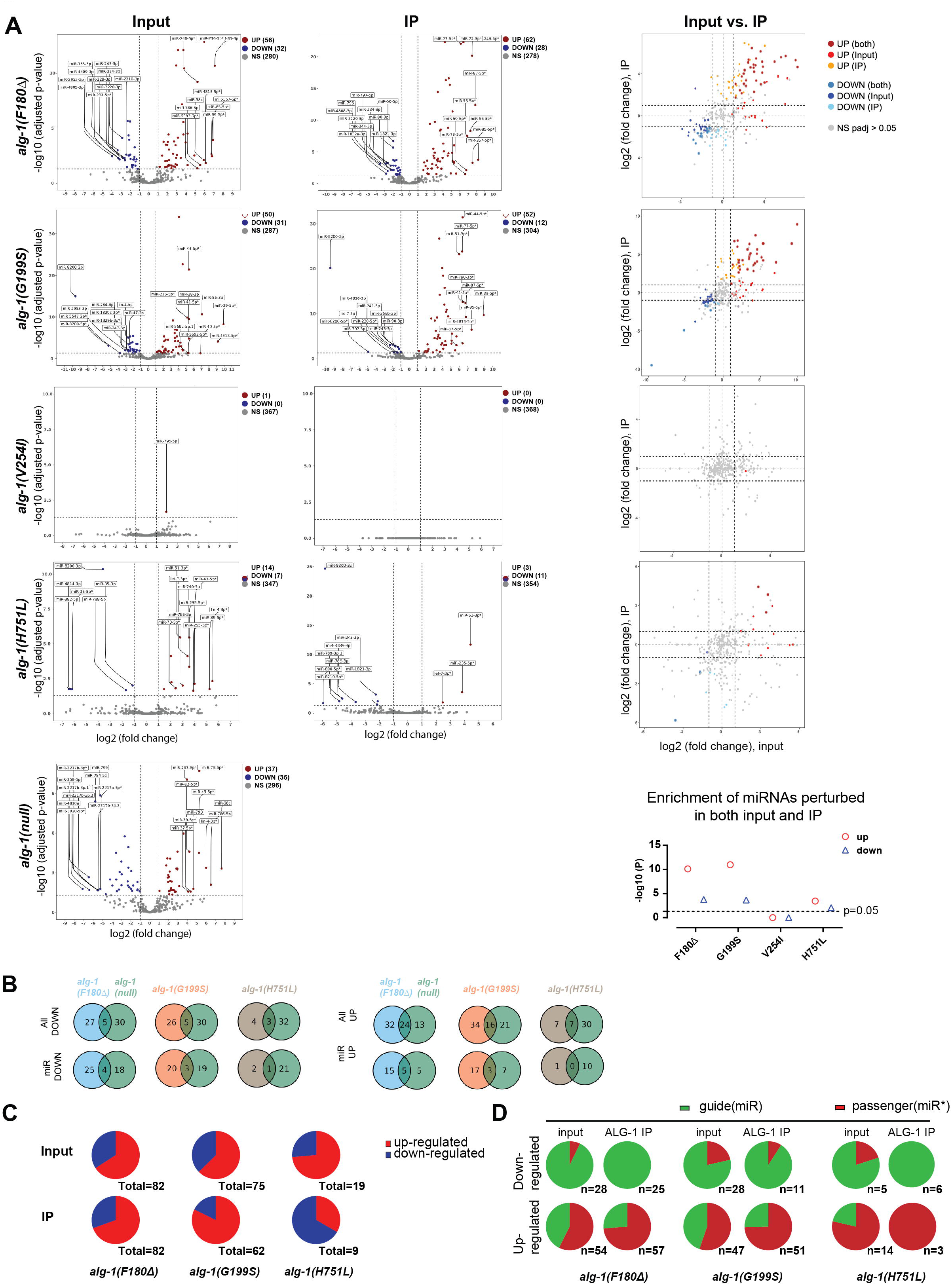
The *alg-1* NDD mutations cause allele-specific disruptions of miRNA expression and miRNA associated with ALG-1 (part 1). **A.** (Left two columns) Volcano plots of normalized miRNA levels in the input and ALG-1 IP. miRNAs with |FC| > 2 and FDR <0.05 are color-coded as red for up-regulation or blue for down-regulation. (Right column) miRNA fold changes comparison between input and ALG-1 IP and the significance of the hypergeometric test of the enrichment of miRNAs that were perturbed in both input and ALG-1 IP (bottom of the column). **B.** Venn diagrams of miRNAs up or down-regulated with statistical significance in the *alg-1(NDD)* and *alg-1(null)* mutants. **C.** Proportions of up/down-regulation among the miRNA perturbation in input and ALG-1 IP. **D.** Proportions of miRNA guide (miR) and passenger (miR*) strands among the perturbed miRNAs.

**Figure S3.**
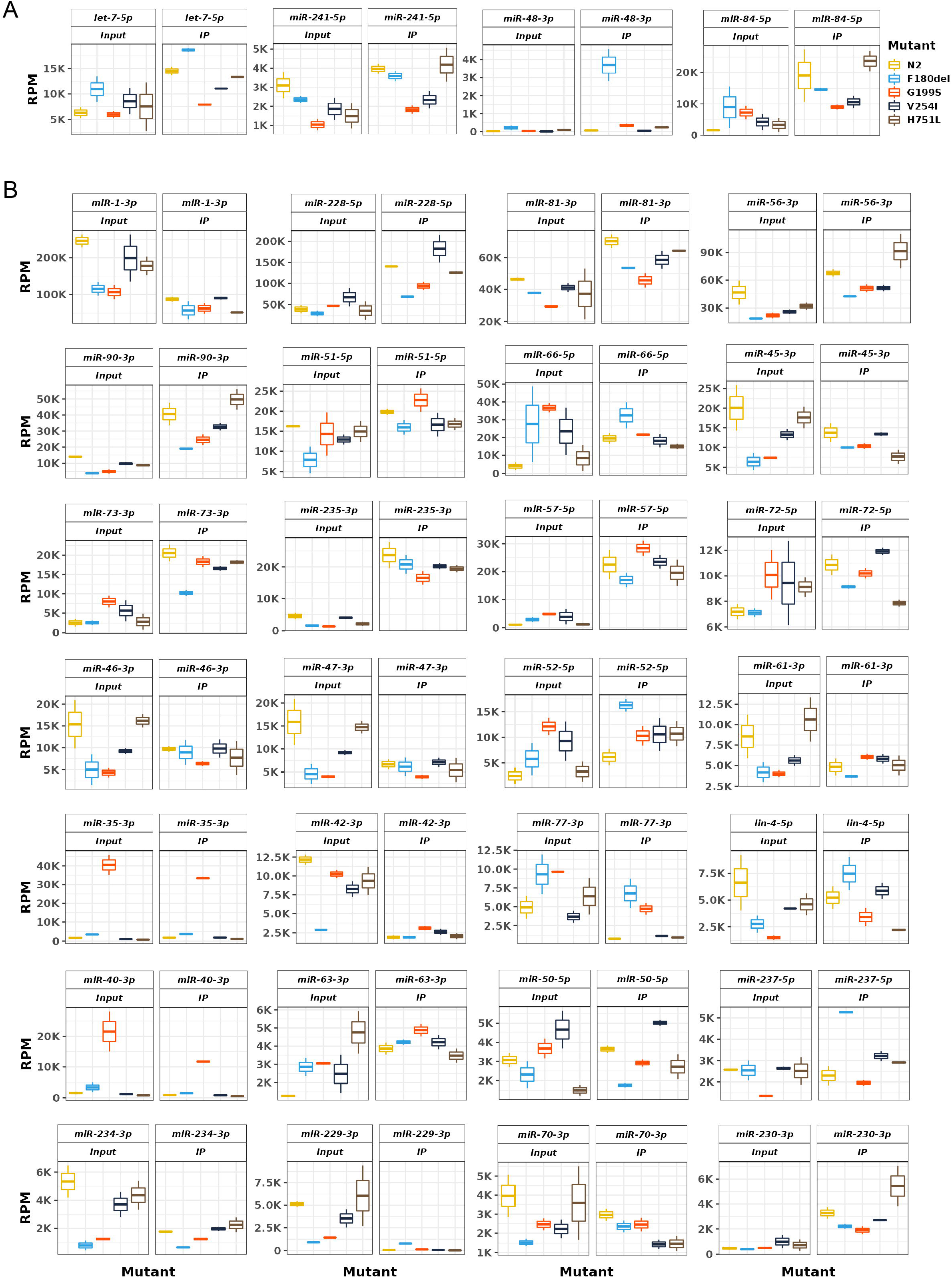
The *alg-1* NDD mutations cause allele-specific disruptions of miRNA expression and/or association with ALG-1 (part 2). **A.** RPM values of major heterochronic miRNAs, whose abundance was altered in either input and/or ALG-1 IP in at least one of the NDD mutants. **B.** RPM values of the most abundant miRNAs with statistically significant abundance (|FC|>2 and p < 0.05) changes in input and/or IP in at least one of the NDD mutants.

**Figure S4.**
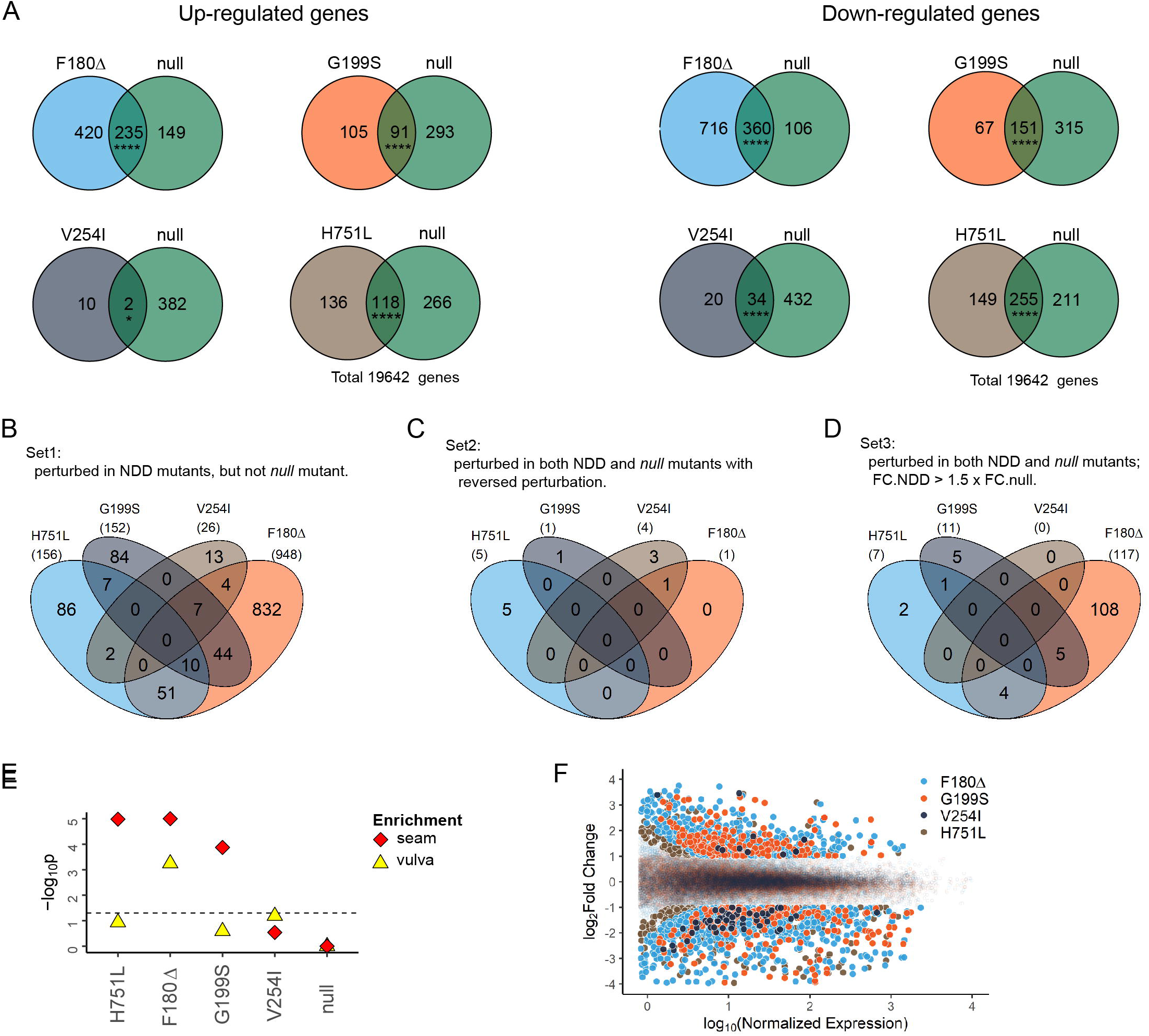
Translatome perturbation profiles between the *alg-1* NDD mutants and the *alg-1 null* mutant partially overlap. **A.** Venn diagrams representing the overlap and distinction of perturbed genes between each *alg-1* NDD mutant and the *alg-1 null* mutant. **B-D.** Venn diagrams for the profile of set1-set3 *amp* genes (Figure 3E) in the *alg-1* NDD mutants. **E.** Enrichment of genes expressed in *C. elegans* vulva and seam cells in the translationally up-regulated genes [41]. Enrichment analysis was tested by a hypergeometric test. ****, p≤0.0001, *, p≤0.05. **F.** MA plots representing the translatome of all the NDD mutants. Solid dots represent perturbed genes with identical statistical significance (|log2FC| > 1.5, p.adj < 0.1).

**Figure S5.**
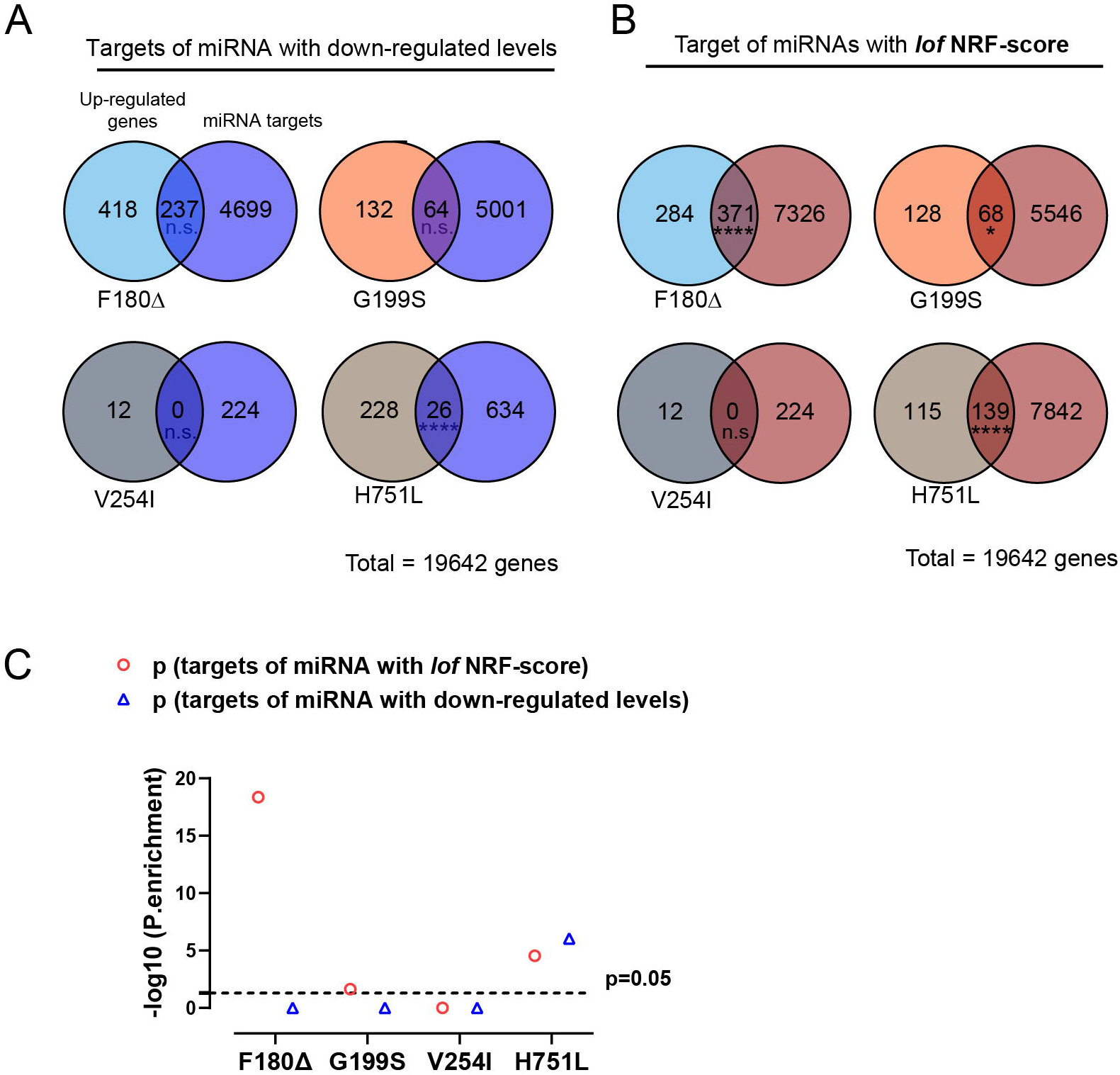
Targets of miRNAs with loss-of-function NRF.score are enriched in the translationally up-regulated genes. **A-B**. Venn diagrams for genes translationally up-regulated in the *alg-1(NDD)* mutants and genes containing target sites for miRNAs with just down-regulated levels (**A)** and with a *lof* NRF.score (**B**). **C.** Significance of hypergeometric test for the enrichment of the targets for miRNAs with decreased expression levels and the targets miRNAs with *lof.* NRF.score in the translationally up-regulated genes.

**Figure S6.**
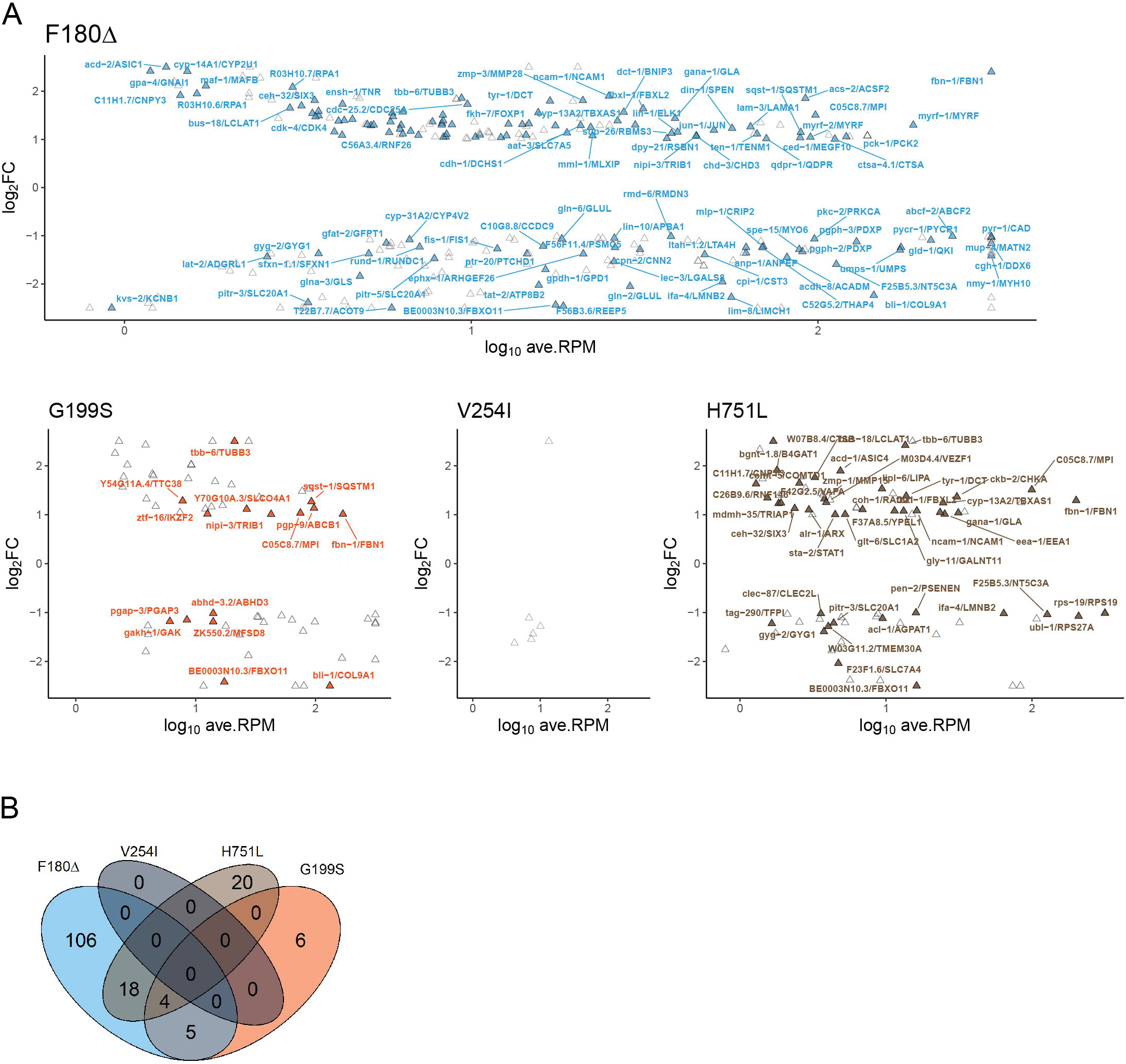
The NDD mutations can perturb the expression of miRNA target genes that have human orthologs with brain expression. **A.** MA plots for the translational levels of up-regulated *C. elegans* genes which have human orthologs with brain translatome expression. Text-labeled and colored points indicate genes that contain target sites of miRNAs with *lof* NRF.score in *C.elegans*. The labels are formatted as Cel_gene_symbol / Hsa_gene_symbol. **B.** Venn diagram for *lof* miRNA (NRF.score < 0.5) target genes that are translationally up-regulated in *C.elegans* and have human ortholog expressed in brain translatome.

**Figure S7.**
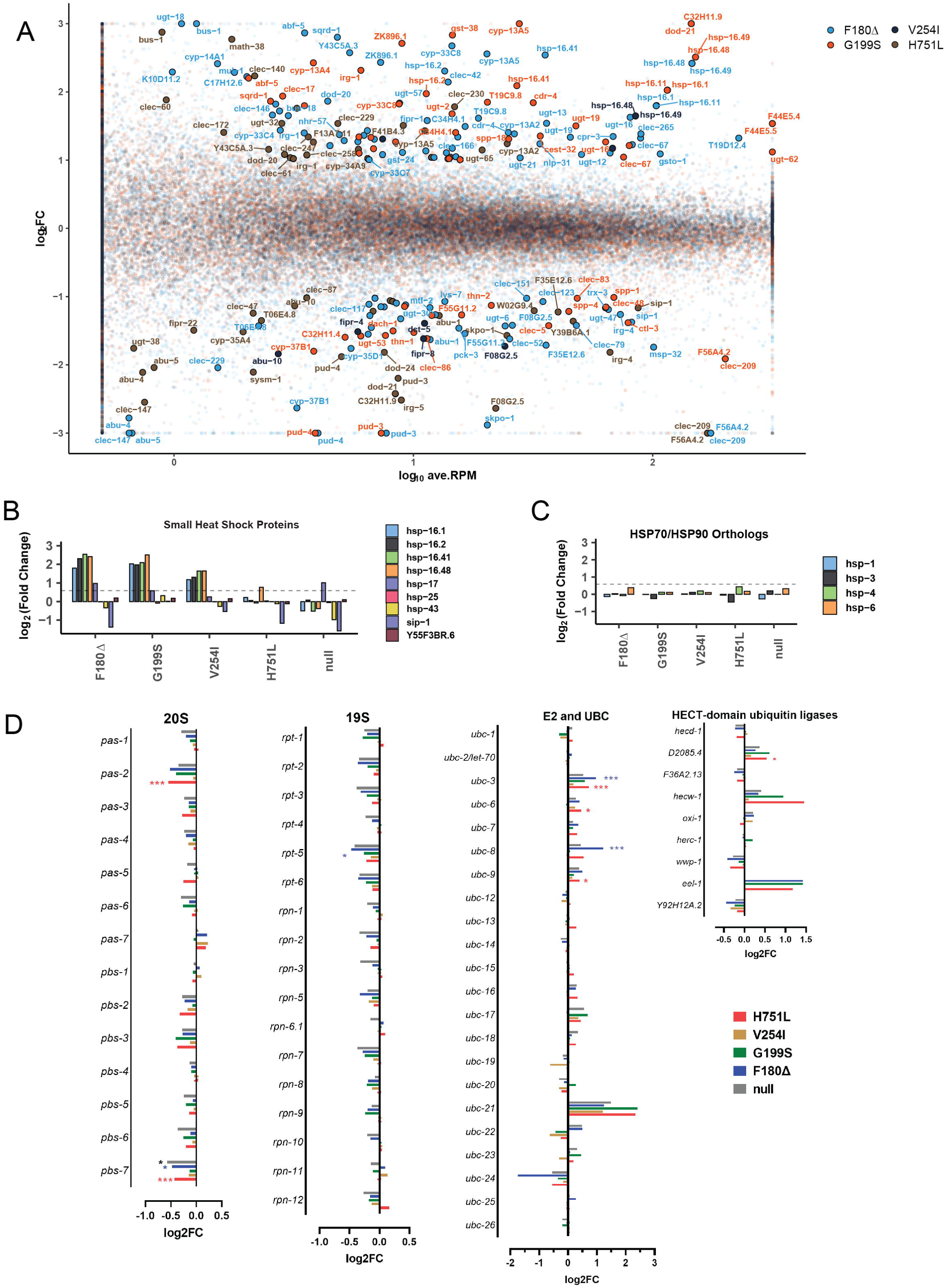
Translational perturbation of stress responses-related genes in the *alg-1* NDD mutant animals. **A.** MA plot of the translational levels of all protein-coding genes. Solid and text-labeled dots represent genes that are related to stress response [50]. **B-C.** Translational levels of small heat shock protein orthologous genes (**B**) and HSP70/HSP90 orthologous genes (**C**) in the NDD mutants and *null* mutants [48]. **D.** RPF levels of the proteasome component proteins and proteolysis-related proteins [77]. Significance tested by DESeq2. ***,p≤0.001; *, p ≤0.05.

**Figure S8.**
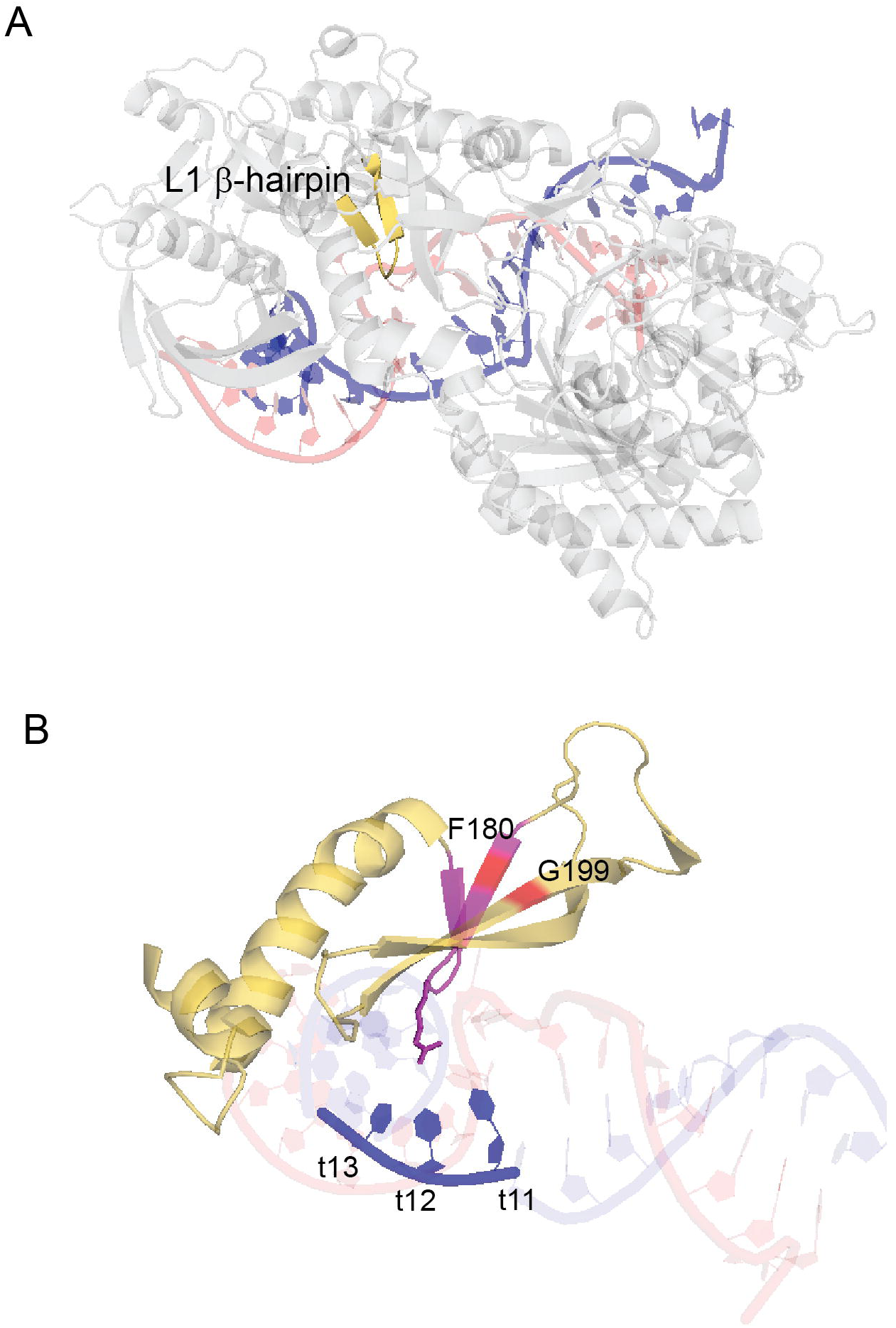
The F180 and G199S may disrupt the function associated with the L1 ß -hairpin. **A.** Visualization of the L1 ß-hairpin (brown) in the *hAGO2*::miRNA::target complex (PDB:: 6MDZ) [31]. Target RNA is colored blue and miRNA is colored red. **B**. Simplified structure of *hAGO2* L1 domain and miRNA::target duplex. L1 ß-hairpin is colored in magenta, with F180 and G199 highlighted in red.

## SUPPLEMENTAL TABLES

**Table S1.Differential expression analyses of total and top changed miRNAs in the alg-1 NDD mutants.** (Related to Figure 3, S2, S3)

**Table S2. Differential expression analyses of ribosome profiling, RNAseq, and translation efficiency (TE) of the alg-1 NDD mutants and analysis of the repressing modes.** (Related to Figure 5, 7, S4)

**Table S3. Lists of antimorphic perturbed (*amp*) genes in the *alg-1* NDD mutants.** (Related to Figure 5, S4)

**Table S4. The net repressive functionality scores (NRF.score) of miRNAs in the alg-1 NDD mutants.**

Only miRNAs with a minimum 15 RPM were analyzed. (Related to Figure 6, S5)

**Table S5. Lists of nervous system-related genes and their differential expression analyses of the translational levels in the alg-1 NDD mutant.**

**A.** Translationally perturbed *C. elegans* genes with orthologs expressed in human brains translatome. **B.** Lists of *C.elegans*/human homologs that were translationally up-regulated in *alg-1* NDD mutants and are related to NDD in sysNDD database (updated to 2.28.2023) [43]. (Related to Figure 8, S6).

**Table S6.**
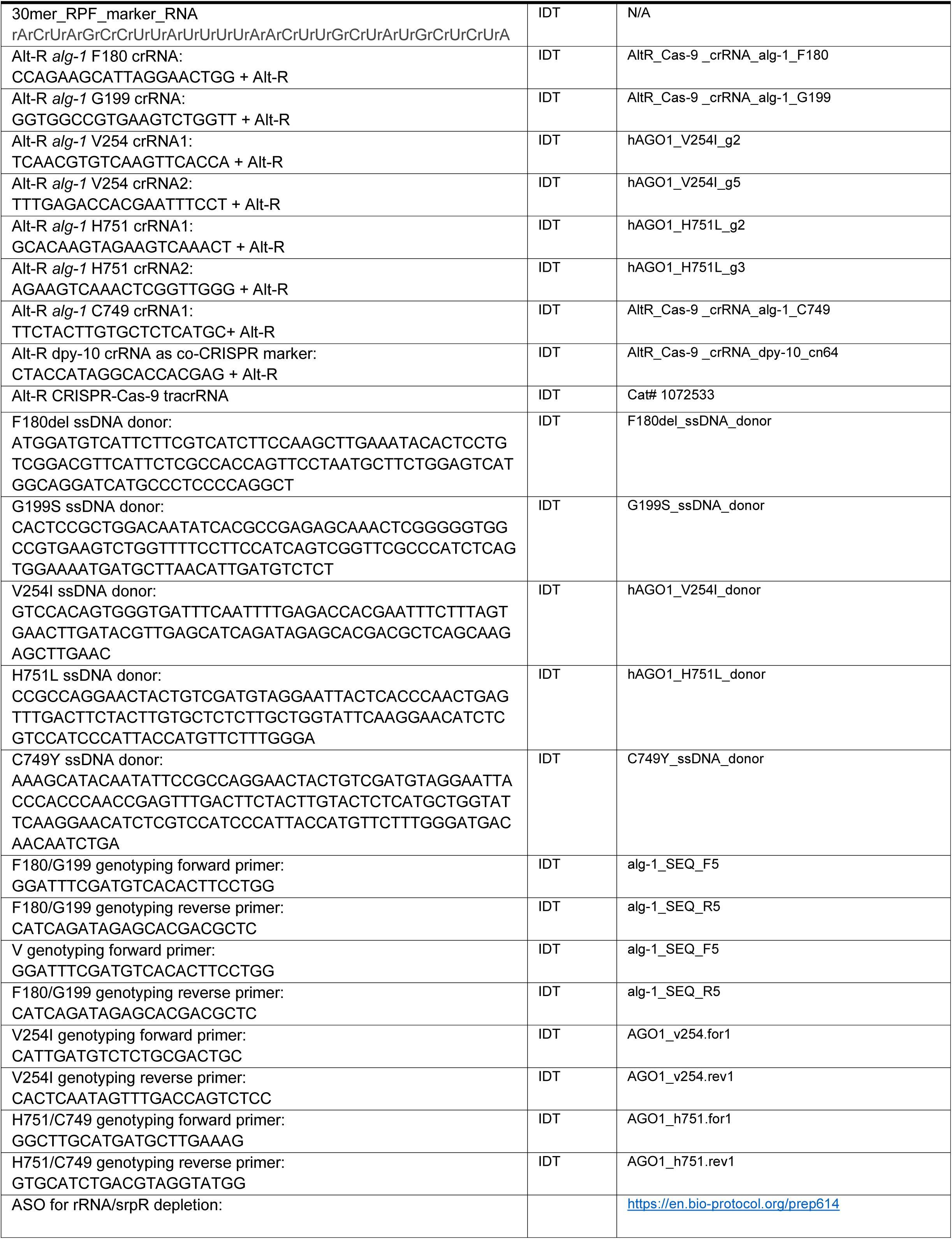
Key oligonucleotides used in this paper for CRISPR/Cas9 mutagenesis.

**Table S7.**
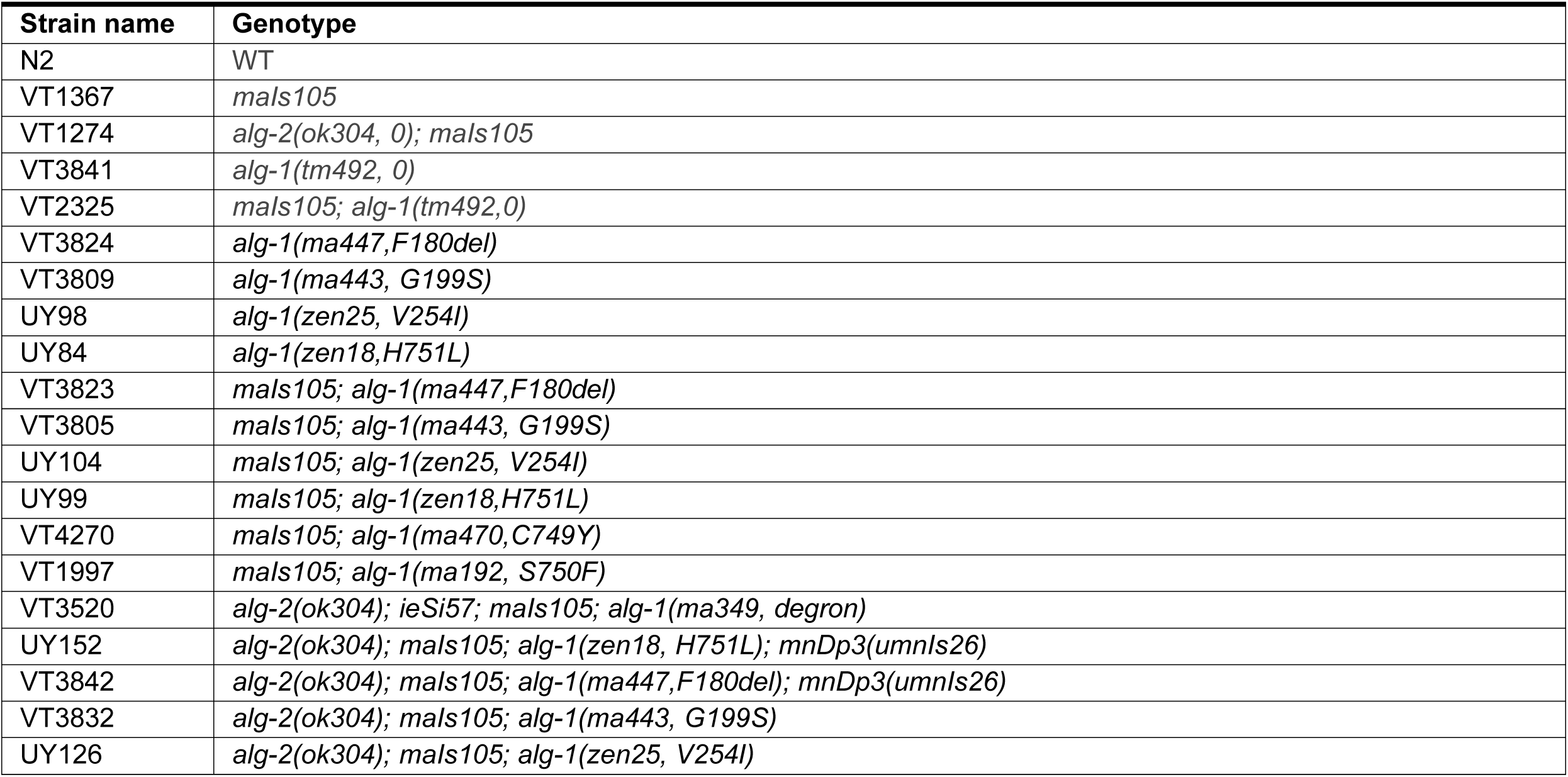
C. elegans strains used in this paper.

## Notes

### Competing Interest Statement

The authors have declared no competing interest.

